# Single-cell longitudinal analysis of SARS-CoV-2 infection in human airway epithelium

**DOI:** 10.1101/2020.05.06.081695

**Authors:** Neal G. Ravindra, Mia Madel Alfajaro, Victor Gasque, Victoria Habet, Jin Wei, Renata B. Filler, Nicholas C. Huston, Han Wan, Klara Szigeti-Buck, Bao Wang, Guilin Wang, Ruth R. Montgomery, Stephanie C. Eisenbarth, Adam Williams, Anna Marie Pyle, Akiko Iwasaki, Tamas L. Horvath, Ellen F. Foxman, Richard W. Pierce, David van Dijk, Craig B. Wilen

## Abstract

SARS-CoV-2, the causative agent of COVID-19, has tragically burdened individuals and institutions around the world. There are currently no approved drugs or vaccines for the treatment or prevention of COVID-19. Enhanced understanding of SARS-CoV-2 infection and pathogenesis is critical for the development of therapeutics. To reveal insight into viral replication, cell tropism, and host-viral interactions of SARS-CoV-2 we performed single-cell RNA sequencing of experimentally infected human bronchial epithelial cells (HBECs) in air-liquid interface cultures over a time-course. This revealed novel polyadenylated viral transcripts and highlighted ciliated cells as a major target of infection, which we confirmed by electron microscopy. Over the course of infection, cell tropism of SARS-CoV-2 expands to other epithelial cell types including basal and club cells. Infection induces cell-intrinsic expression of type I and type III IFNs and IL6 but not IL1. This results in expression of interferon-stimulated genes in both infected and bystander cells. We observe similar gene expression changes from a COVID-19 patient *ex vivo*. In addition, we developed a new computational method termed CONditional DENSity Embedding (CONDENSE) to characterize and compare temporal gene dynamics in response to infection, which revealed genes relating to endothelin, angiogenesis, interferon, and inflammation-causing signaling pathways. In this study, we conducted an in-depth analysis of SARS-CoV-2 infection in HBECs and a COVID-19 patient and revealed genes, cell types, and cell state changes associated with infection.

## 1 Introduction

In December 2019, a novel viral pneumonia, now referred to as Coronavirus Disease 2019 (COVID-19), was observed in Wuhan, China [1]. Severe Acute Respiratory Syndrome (SARS)-Coronavirus (CoV)-2, the causative agent of COVID-19, has caused a staggering and growing number of infections and deaths globally. There are currently no approved drugs or vaccines for the treatment or prevention of COVID-19. Clinical presentation is highly variable ranging from asymptomatic infection to acute respiratory distress syndrome and death [2]. Enhanced understanding of viral pathogenesis at the cellular and molecular level is critical for development of prognostic tools and novel therapeutics.

CoVs are enveloped viruses with positive-sense, single-stranded RNA genomes ranging from 26–30 kb [3]. Six human CoVs have been previously identified: HCoV-NL63 and HCoV-229E, which belong to the Alphacoronavirus genus; and HCoV-OC43, HCoV-HKU1, SARS-CoV, and Middle East Respiratory Syndrome CoV (MERS-CoV), which belong to the Betacoronavirus genus [4]. In the past two decades, CoVs have become a major public health concern due to potential zoonotic transmission, as revealed by the emergence of SARS-CoV in 2002, which infected 8, 000 people worldwide with a mortality rate of 10–15%, and MERS-CoV in 2012 and 2019, which infected 2, 500 people with a mortality rate of 35%, and now SARS-CoV-2 [5]. Tissue and cell tropism are key determinants of viral pathogenesis. SARS-CoV-2 entry into cells depends on the binding of the viral spike (S) protein to its cognate receptor angiotensin-converting enzyme II (ACE2) on the cell surface [2]. ACE2 is also the receptor for SARS-CoV and HCoV-NL63, yet these viruses induce profoundly different morbidity and mortality suggesting unknown determinants of coronavirus pathogenesis [6, 7]. Additionally, proteolytic priming of the S protein by host proteases is also critical for viral entry [8]. The cellular serine protease Type II transmembrane (TMPRSS2) is used by SARS-CoV-2 for S protein priming [9, 8, 10, 11]. This is also used by SARS-CoV alongside the endosomal cysteine proteases cathepsin B and L [12, 13]. Another host protease, furin, has been suggested to mediate SARS-CoV-2 pathogenesis; however, the precise role of host proteases in SARS-CoV-2 entry remains to be determined [14, 11].

SARS-CoV and MERS-CoV caused fatal pneumonia associated with rapid virus replication, elevation of proinflammatory cytokines, and immune cell infiltration [15]. These characteristics are similarly observed in SARS-CoV-2 infection. COVID-19 patients have increased levels of proinflammatory effector cytokines, such as TNF*α*, IL1B, and IL6, as well as chemokines, such as CCL2 and CXCL10, especially in those who are critically ill [16, 17, 18, 19]. These studies suggest that an over exuberant immune response characterized by cytokine storm rather than direct virus-induced damage may be responsible for COVID-19 pathogenesis. The cell types and mechanisms underlying this immune response are unclear for SARS-CoV-2. In addition, it has been observed that age is a strong risk factor for more severe disease (CDC MMWR). In the United States, between February 12 and March 16, 2020, the case-fatality rate was 10.4 − − 27.3% for patients ≥ 85 years old, compared to 0.1 − − 0.2% for patients 20 − − 44 years old (CDC MMWR). The reason for this increased risk remains unknown.

Our knowledge of SARS-CoV-2 biology and pathogenesis is incomplete. To address this gap, we performed single-cell (sc) RNA sequencing (RNA-seq) on organotypic human bronchial epithelial cells (HBECs) infected with SARS-CoV-2. This culture system supports epithelial cell differentiation and mimics key aspects of the mucosal epithelium. By utilizing scRNA-seq and electron microscopy, we revealed that ciliated cells are a major target of SARS-CoV-2 infection. During the course of infection, cell tropism of SARS-CoV-2 extended to other epithelial cells including basal and club cells. Furthermore, SARS-CoV-2 infection elicited intrinsic expression of type I and type III interferons and IL6 but not IL1. Interferon stimulated gene (ISG) expression was observed in both infected and bystander cell populations. We also found that differentially expressed genes between infected and bystander ciliated cells were enriched for age-associated genes, a critical risk factor for COVID-19. Here, we provide a detailed analysis of SARS-CoV-2 infection in HBECs and in cells from a pediatric COVID-19 patient during peak and resolving disease. We revealed novel SARS-CoV-2 transcripts, the cell tropism, host gene expression and cell state related to infection.

## 2 Results

### 2.1 Viral infection dynamics

To characterize SARS-CoV-2 interactions with the human airway, we performed single-cell RNA sequencing of SARS-CoV-2 infected airway epithelium. We cultured primary HBECs at an air-liquid interface (ALI) for 28 days and then challenged the apical surface of the epithelium with 10^4^ plaque forming units (PFU) of SARS-CoV-2 (Fig 1A). Exponential viral replication over the course of the experiment was demonstrated by qRT-PCR of cell lysate for the SARS-CoV-2 nucleocapsid (N) gene (Fig 1B). At 1, 2, and 3 days post-infection (dpi), a single cell suspension was generated and 3’ single-cell RNA sequencing was performed on 77,143 cells across four samples cells per sample with an average of 31,383 reads per cell (Fig 1C, S1A). To define SARS-CoV-2 infected cells, we mapped reads to the viral reference genome and quantified viral transcript abundance on a per cell basis (Fig 1D). We defined productively infected cells as those with at least ten viral transcripts per cell, which controls for background due to misaligned reads in the mock sample. Consistent with viral genome replication (Fig 1B), we observed a time-dependent increase in the abundance of infected cells from 1 to 3 dpi (Fig 1E, S1B).

**Figure 1:**
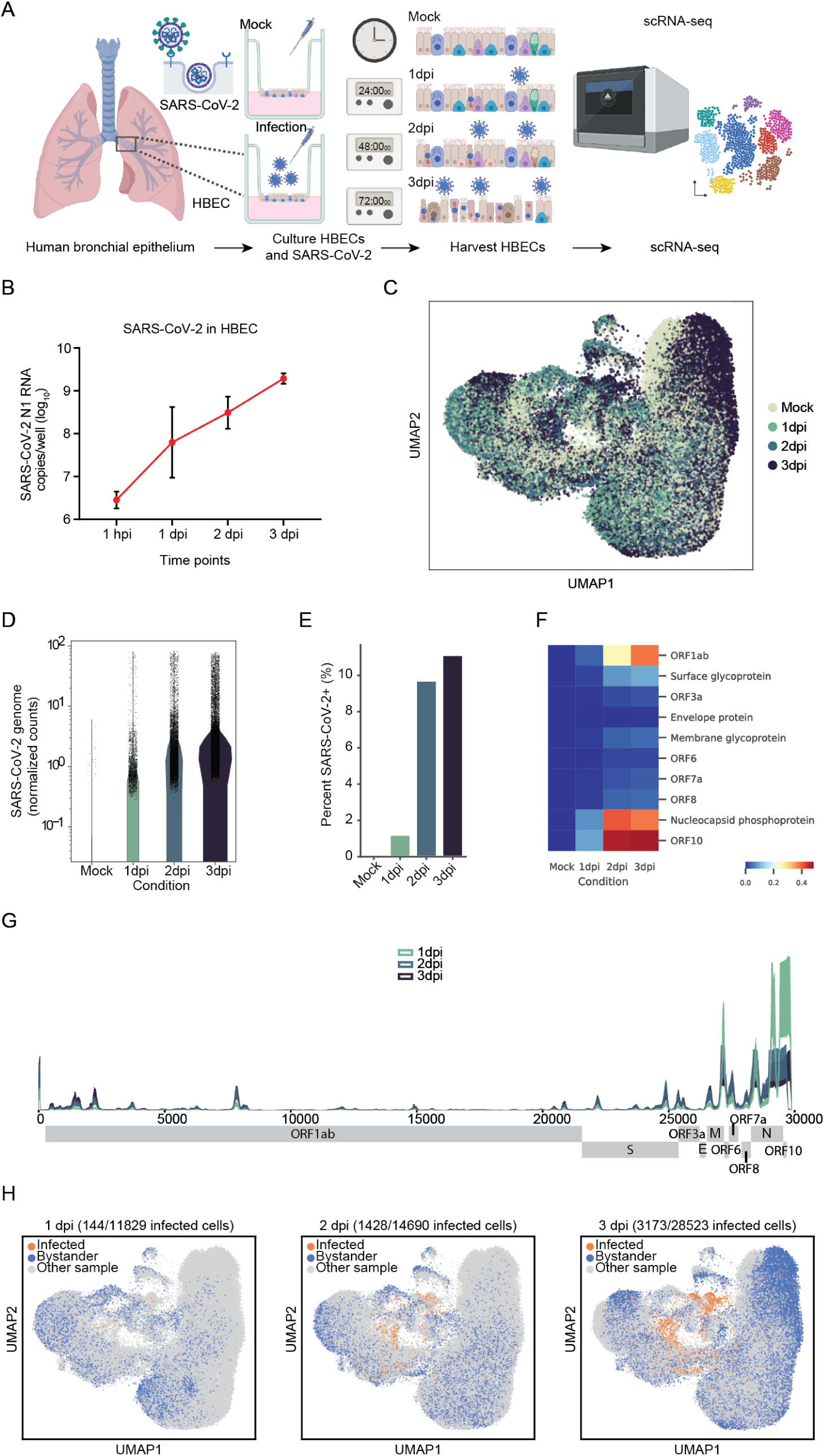
scRNA-seq reveals SARS-CoV-2 infection of HBECs. **A.** Schematic of the experiment. Human bronchial epithelial cells (HBECs) were cultured and infected or not (mock) with SARS-CoV-2. Infected cultures were collected for scRNA-seq at 1, 2 and 3 days post infection (dpi). **B.** RT-qPCR in cultured HBEC to detect viral transcripts at 1 hour post-infection (hpi), 1, 2, and 3 days post-infection (dpi) (copies/well). **C**. UMAP visualization of the scRNA-seq gene counts after batch correction. Each point represents a cell, colored by sample. **D.** Normalized counts of viral counts in each condition. For each cell, viral counts were determined by aligning reads to a single, genome-wide reference. **E.** Percent of cells infected by SARS-CoV-2, based on a viral genes count threshold (see Materials and Methods) **F.** Normalized heatmap of the viral Open Reading Frame (ORF) counts in each condition. Reads were aligned to each 10 SARS-CoV-2 ORFs. **G.** Coverage plot of viral reads aligned to SARS-CoV-2 genome. The sequencing depth was computed for each genomic position for each condition. As infection progresses, coverage becomes more dispersed on the genome. **H.** UMAP visualizations of infected and bystander cells in each condition after batch correction. Bystander cells are defined as cells that remained uninfected in HBEC samples challenged with SARS-CoV-2.

Next, we characterized the SARS-CoV-2 transcriptome at the single-cell level. In addition to the reads expected to align immediately upstream of the canonical SARS-CoV-2 poly-A tail, our results show additional reads aligning elsewhere in the genome suggesting the existence of non-canonical, poly-adenylated sub-genomic RNAs (sgRNAs) (Fig 1F, G). The distribution of polyadenylated viral transcripts shifts from 3’ to 5’ during the infection time-course (Fig 1F,G,S1C). Mapping of productive infected cells reveals multiple infected cell clusters that expand over time and are not present in the mock sample (Fig 1H). Using RT-PCR, we successfully validate two unique peaks, one that mapped in the middle of the Open Reading Frame (ORF)1ab region, and a second peak that mapped near the ORF6 boundary (Fig S1D, top panel). Our results confirm that RT-PCR products corresponding to each of the two peaks appears after 2 dpi (Fig S1D, bottom panel, red arrows). Importantly, the absence of these RT-PCR bands in the mock and 1 dpi samples suggests they are not the result of non-specific oligo-d(T) priming of cellular or viral RNAs. We included two positive controls, amplifying RT-PCR products of increasing length from the canonical SARS-CoV-2 poly-A tail (Fig S1D, bottom panel, green arrows). These RT-PCR bands appear as early as 1 dpi, are specific to infected cells and run at their expected lengths. These positive controls validate that we are able to capture known poly-adenylated viral transcripts with this RT-PCR priming strategy.

### 2.2 Identification of the cell tropism of SARS-CoV-2

The human airway is comprised of diverse epithelial cell types with critical functions in gas exchange, structure, and immunity. We sought to determine the cellular tropism of SARS-CoV-2 in the bronchial epithelium, as the airway is a critical target of viral pathogenesis. We identified eight major clusters comprising canonical epithelial cell types: ciliated cells, basal cells, club cells, goblet cells, neuroendocrine cells, ionocytes, and tuft cells (Fig 2A, S2A). We also observed a cell population intermediate between basal cells and club cells (BC/club) likely representing basal stem cells differentiating into club cells. Analysis of differentially expressed genes show these cell clusters express classical epithelial cell type-specific markers (Fig 2B). Mapping viral infected cells within specified epithelial cell types reveals that ciliated, basal, club, and BC/Club cells are susceptible to SARS-CoV-2 infection whereas goblet, neuroendocrine, tuft cells, and ionocytes are relatively resistant to infection (Fig 2C,D). At 1 dpi, ciliated cells represent 83% of infected cells and continue to comprise the majority of infected cells throughout infection (Fig 2E,F). However, during productive infection, the number of infected basal, club, and BC/club cells also increases, suggesting that these cells are major secondary targets (Fig 2E,F). The distribution of polyadenylated viral transcripts along the length of the genome is similar across infected cell types (Fig S2C).

**Figure 2:**
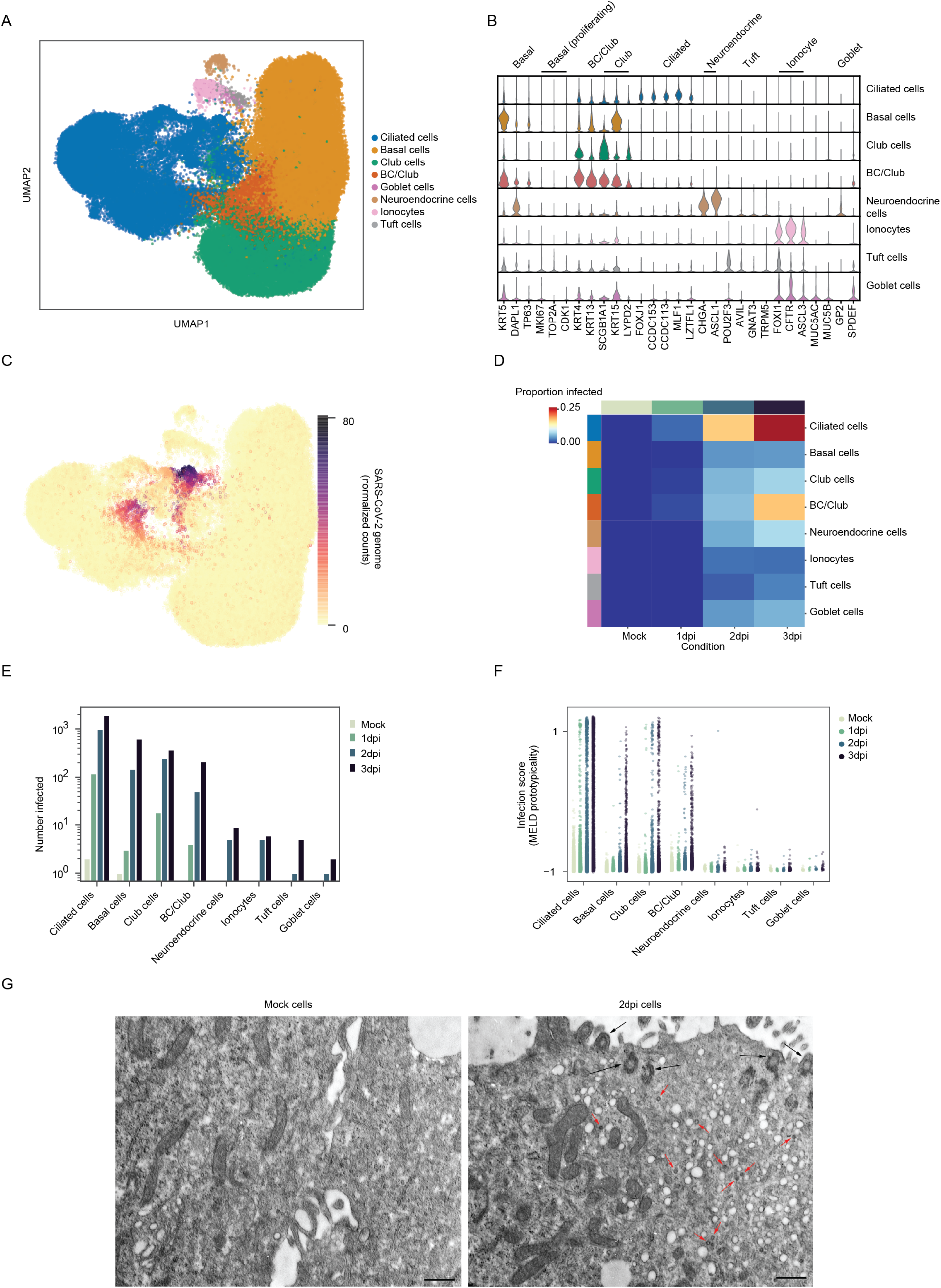
SARS-CoV-2 cell tropism. **A.** UMAP visualization of the cell clusters manually annotated. Cells were first clustered with the Louvain algorithm, then annotated according to a panel of marker genes. **B.** Violin plot of annotation marker genes and SARS-CoV-2 putative relevant genes based on the recent literature. **C.** Uniform Manifold Approximation and Projection (UMAP) visualization of the normalized counts of SARS-CoV-2 reads. Reads were determined here as in Figure 1D.**D.** Proportion of infected cells across conditions and cell types. **E.** Histogram of the number of infected cells per cell type across conditions. **F.** Infection score inferred from Manifold Enhancement of Latent Dimensions (MELD) showing extent of infection per cell stratified by cell time. **G.** Transmission electron microscopy image of mock (left) and SARS-CoV-2 human bronchial epithelial cell (HBEC) reveal infected ciliated cells at 2 days post infection (dpi) (right). Scale (bottom) corresponds to 500 nm. Red arrows denoted virus particles and black arrows cilia.

To independently verify SARS-CoV-2 cell tropism, HBECs cultured under identical conditions as for scRNA-seq were assessed by transmission electron microscopy. At 2 dpi, we observed numerous virus particles approximately 80 nm in size in ciliated cells (Fig 2G). This is consistent with the known size distribution of coronaviruses [20]. These particles were not observed in a mock control sample (Fig 2G). Together, this confirms that ciliated cells are a major target of SARS-CoV-2 infection in the human bronchus.

### 2.3 Determinants of cell tropism

Next, we sought to determine the host transcriptional correlates of SARS-CoV-2 cell tropism. As viral entry is a major determinant of cell tropism, we first investigated whether expression of the SARS-CoV-2 receptor ACE2 predicted infection. We observed ACE2 expression at low levels across ciliated, basal, club and BC/club cells in the mock condition (Fig 3). Surprisingly, ACE2 expression was poorly correlated with SARS-CoV-2 infection on a per cell basis (Spearman’s r between viral genome and ACE2 in ciliated cells, -0.06, and between ACE2 and infection score in ciliated cells, 0.09). However, ACE2 expression was increased in the four susceptible cell populations: ciliated, basal, club, and BC/club relative to the non-susceptible cell types: neuroendocrine, ionocytes, goblet, and tuft cells (Fig S3). Expression of ACE and CLTRN which are structural homologs of ACE2 and the aminopeptideases ANPEP and DPP4 (the MERS-CoV receptor) were also poorly correlated with SARS-CoV-2 susceptibility (Fig 3B-E).

**Figure 3:**
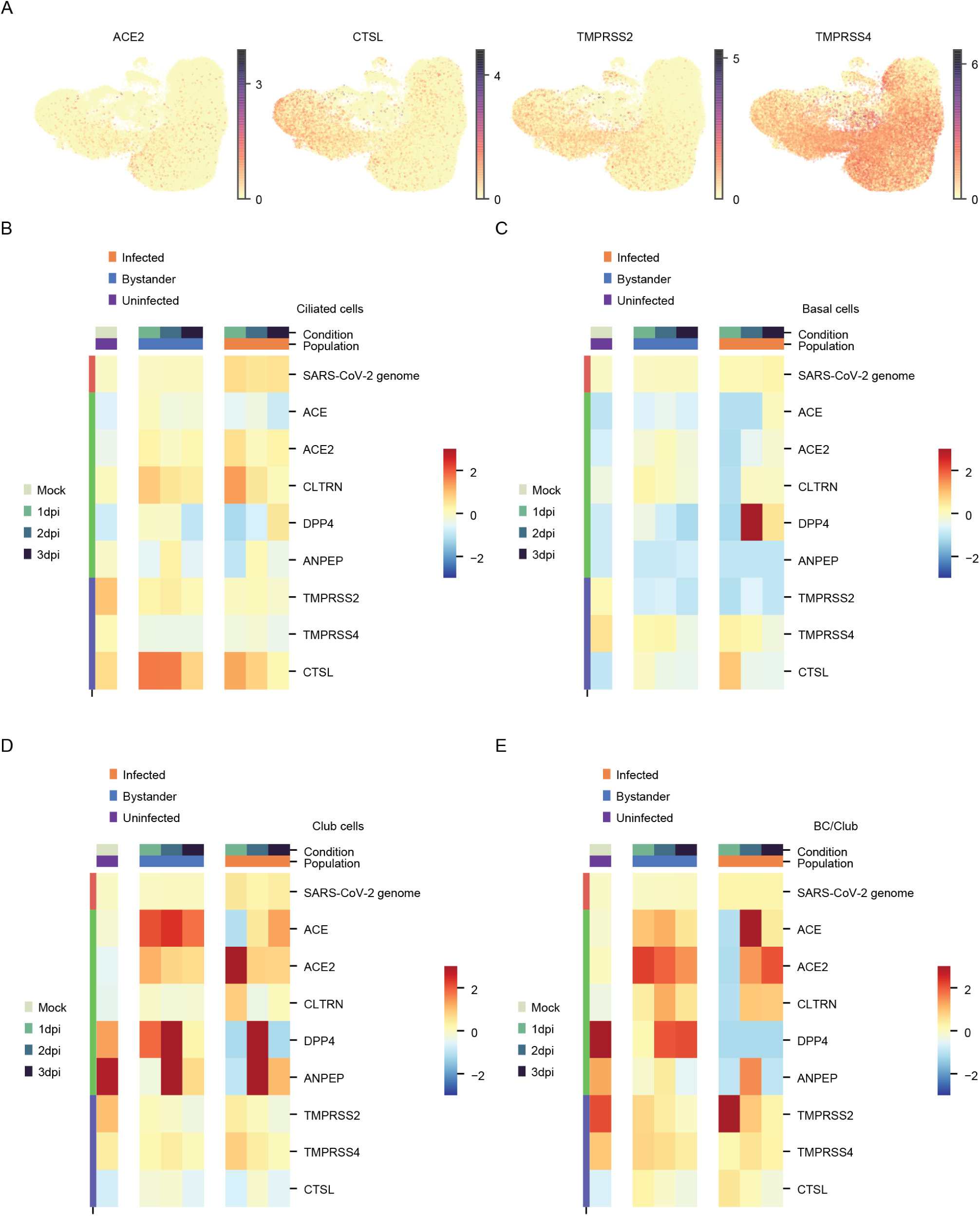
Expression of known entry determinants across bronchial epithelial cell types. **A.** UMAP visualizations, colored by expression of four receptors and proteases expressed in human bronchial epithelial cells (HBECs): ACE2, TMPRSS2, TMPRSS4 and CTSL. **B-E.** Heatmaps of receptors and proteases in ciliated (**B.**), basal (**C.**), club (**D.**) and BC/Club cells (**E.**).

ACE2 was recently demonstrated to be an interferon-stimulated gene (ISG) [21]. Here we observe a modest increase in ACE2 expression in both infected and bystander cells relative to the mock sample consistent with dynamic regulation of ACE2 expression by the host innate immune response to SARS-CoV-2 (Fig 3B-E). To examine whether expression of other potentially pro-viral genes explain SARS-CoV-2 cell tropism, we assessed the expression of the proteases that may potentiate SARS-CoV-2 infection. The trans-membrane serine protease TMPRSS2 and cathepsin L have been implicated in SARS-CoV-2 entry [22]. We also examined the related protease TMPRSS4 which cleaves influenza hemaglutinin, similar to TMPRSS2, and may also play a role in SARS-CoV-2 entry [22, 23]. TMPRSS2 and CTSL were expressed predominantly in basal, club and ciliated cells while TMPRSS4 was broadly expressed in all epithelial cell types (Fig 3B-E). The specific role of proteases in governing SARS-CoV-2 tropism in the human airway epithelium remains to be further elucidated.

### 2.4 Innate immune response to SARS-CoV-2 infection

SWe investigated the host transcriptome to assess the host immune response to SARS-CoV-2 infection at single-cell resolution in the human airway epithelium. We observed robust induction of both type I interferon (IFNB1) and type III interferons (IFNL1, IFNL2, and IFNL3) in ciliated, basal, club, and BC/club cells co-expressing SARS-CoV-2 transcripts (Fig 4). Interestingly, the kinetics of IFNB1 induction were delayed relative to type III interferon. In contrast, there was scant IFN induction in uninfected ciliated, basal, club, and BC/club cells. We also did not observe IFN induction in neuroendocrine, ionocytes, goblet, or tuft cells consistent with these cell types not being major target cells of SARS-CoV-2 (Fig S4). This demonstrates direct SARS-CoV-2 infection of a given cell is critical for interferon induction. Type I and III interferons signal through IFNAR and IFNLR, respectively, resulting in expression of hundreds of ISGs. Consistent with this, we observed broad ISG induction (IFI27, IFITM3, IFI6, MX1, and ISG15) in both infected and bystander cells of all cell types (Fig 4, S4) suggesting IFN from infected cells is acting *in trans* on both infected cells and uninfected bystander cells.

**Figure 4:**
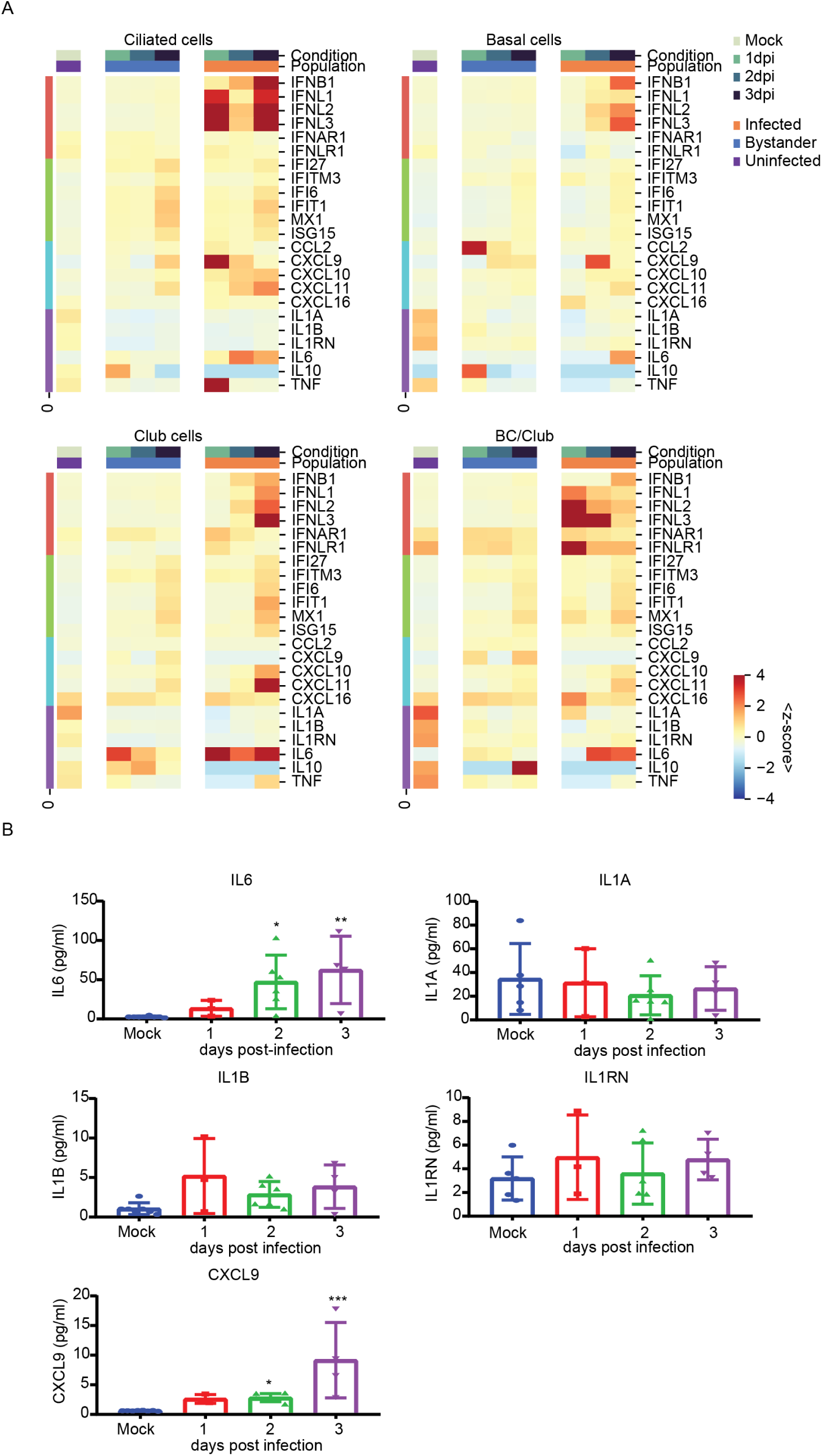
SARS-CoV-2 infection induces robust innate immune response. **A-D.** Heatmaps of expression of key innate immune and inflammatory genes in ciliated (**A.**), basal (**B.**), club (**C.**) and BC/club cells (**D.**) This reveals infected cells up-regulate type I and III interferons, IL-6, and chemokines in a cell-intrinsic manner while interferon-stimulated genes are induced in both infected and bystander cells. Infection also stimulates IL-6 protein secretion. Minimal changes in IL-1A, IL-1B, and IL1RN expression and protein secretion are observed.

The host anti-viral response also results in chemokine induction leading to recruitment of immune cells, a hallmark of severe COVID-19 [24]. Here, we observe induction of CXCL9, CXCL10, and CXCL11 which propagate signals through the cognate CXCR3 receptor to recruit activated T cells and NK cells (Fig 4). This induction was evident in infected but not bystander cells (Fig 4). In contrast, CCL2 and CXCL16 which recruit monocytes and T cells, respectively, were not dynamically regulated over the conditions evaluated (Fig 4 and S4). We also observed substantial induction of the pro-inflammatory cytokine IL-6 in infected ciliated, basal, club, and BC/club cells but not in uninfected bystander cells of these same populations. Interestingly, expression of pro-inflammatory IL-1 was modestly downregulated in all cell types after infection whereas IL-10 and TNF*α* expression were not significantly regulated by infection in this system (Fig 4). We further characterized the levels of selected proteins in the basolateral supernatant of mock and infected HBECs and observed induction of IL6 and CXCL9 but not IL1A, IL1B, and IL1RN, consistent with gene expression changes (Fig 4E).

### 2.5 Differentially expressed genes in response to SARS-CoV-2 infection

To determine how SARS-CoV-2 infection perturbed the cellular transcriptome, we computationally pooled the three infected samples and analyzed the top 100 differentially expressed genes between infected and uninfected bystander cells of a given cell type within the 1, 2, and 3 dpi samples (Fig 5A). PANTHER gene ontology analysis revealed infected ciliated cells had increased expression of genes involved in apoptosis (e.g. PMAIP1, SQSTM1, ATF3), translation initiation and viral gene expression (e.g. RPS12, RPL37A) and inflammation (e.g. NFKBIA and NFKBIZ) compared to bystander cells (Fig 5B,C and differentially expressed gene lists in Supplemental Files). Similar genes are enriched in other infected cell populations (Fig S5). In contrast, infected ciliated cells showed significant downregulation of genes included in biological processes involved in cilium function (e.g. DYNLL1), calcium signaling (e.g. CALM1, CALM2), and iron homeostasis (e.g. FTH1, FTL; Fig 5B,C and S5).

**Figure 5:**
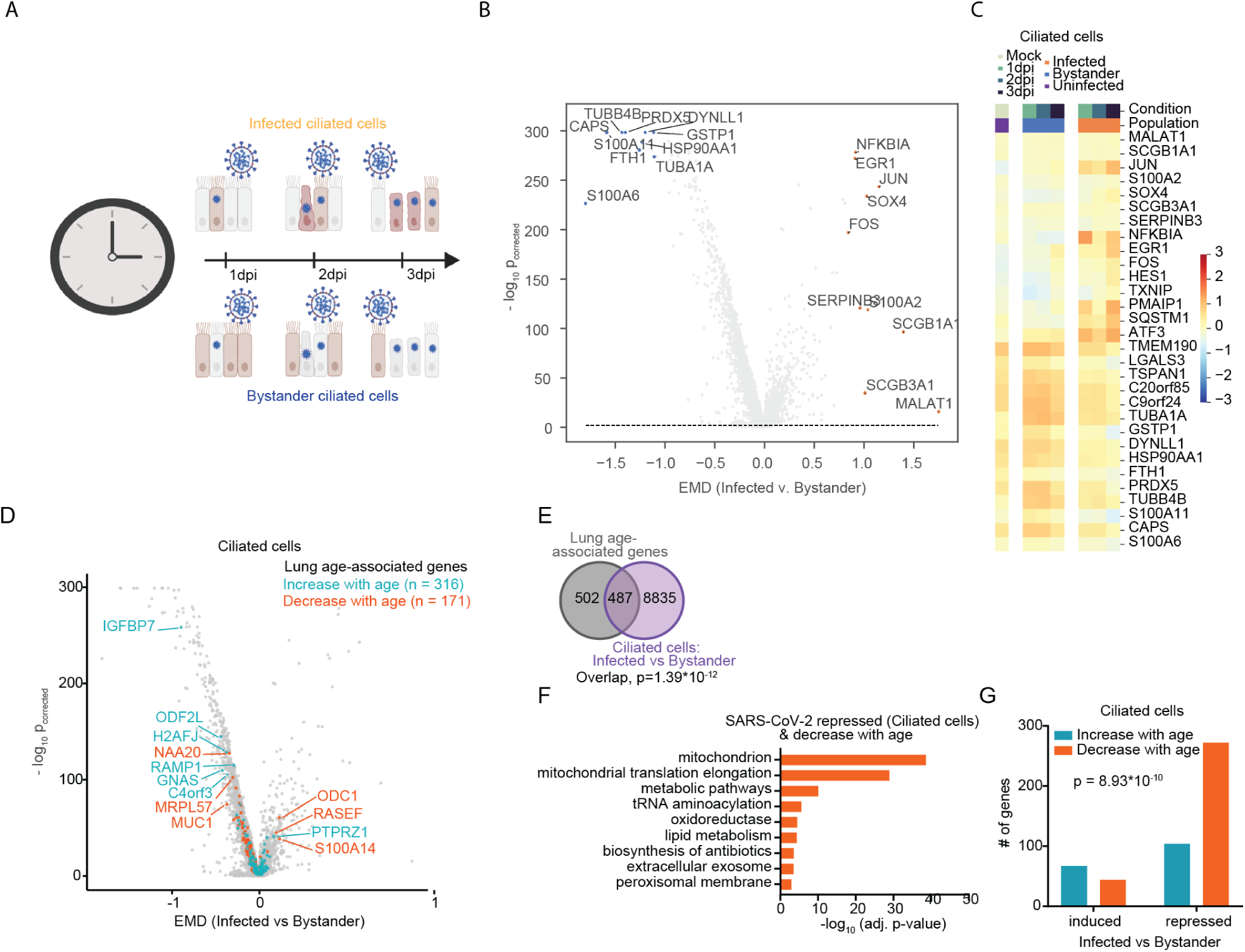
Expression of differentially expressed genes. (**A**) Schematic of the differential expression analysis. Two main cell populations are observed: bystander cells that were not infected by the virus at 3 days post infection (dpi) and infected cells that contain active viral replication and transcription at 3 dpi. (**B**) Volcano plots highlighting the most differentially expressed genes between infected and bystander cells in ciliated cells at 3 dpi as measured by earth mover’s distance (EMD). (**C**) Heatmap of the most differentially expressed genes between uninfected, infected and bystander cells in ciliated cells in all conditions. (**D**) Volcano plot of differentially expressed genes in ciliated cells (infected vs. bystander). Age-associated genes in human lung are color-coded (blue, increase in expression with age; orange, decrease in expression with age. (**E**) Venn diagram highlighting the intersection between lung age-associated genes and SARS-CoV-2 regulated genes in ciliated cells. Statistical significance of the overlap was assessed by hypergeometric test. (**F**) Characteristics of age-associated genes that are affected by SARS-CoV-2 infection in ciliated cells. Statistical significance of the interaction between the directionality of SARS-CoV-2 regulation (induced or repressed) and the directionality of age-association (increase or decrease with age) was assessed by two-tailed Fischer’s exact test. (**G**) DAVID gene ontology and pathway analysis of genes repressed by SARS-CoV-2 infection in ciliated cells that also decrease in expression with aging.

Given that age is a major risk factor for COVID-19 severity, we explored whether the expression of these differentially expressed genes may naturally change with age [25]. We analyzed the differentially expressed genes between infected and bystander ciliated cells for previously determined age-associated genes (Figure 5D) [26]. Age-associated genes were significantly enriched among the differently expressed genes (Figure 5E). Of these genes at the intersection, genes repressed by SARS-CoV-2 infection tended to decrease in expression with age, and vice versa (Figure 5G). Analysis of the genes that are repressed by SARS-CoV-2 infection and that decrease in expression with age demonstrated enrichment for several gene ontologies and pathways, most notably mitochondria-related pathways (Figure 5F). Together this suggests that SARS-CoV-2 infection reprograms the cellular transcriptome, resulting in promotion of viral infection and potentially resulting in cell dysfunction and apoptosis that contributes to COVID-19 pathogenesis.

### 2.6 Gene dynamics in infected and bystander cells using CONDENSE

To investigate transcriptional dynamics in response to SARS-CoV-2 infection, we developed a new computational method for characterizing genes by their temporal dynamics (Methods, Figure 6A) termed CONditional DENSity Embedding (or CONDENSE). Briefly, CONDENSE uses a latent (or pseudo) time dimension and conditional kernel density estimation to quantify, per gene, the dynamics along the time axis. In the current application this is done separately for infected and bystander cells. CONDENSE then computes distances between the density estimates providing a metric space that can be used for manifold learning. The result is an embedding of the genes according to their dynamics that can subsequently be used for down stream analyses such as visualization, clustering, and archetypal analysis. Here, we use archetypal analysis to characterize the gene dynamics manifold and observe that, between infected and bystander cells, genes may exhibit similar dynamics over time (e.g., Archetypes 7) while others exhibit differential dynamics (e.g., Archetypes 1, 3, 7, 18) (Figure 6B,S6). Individual genes can be represented as a combination of these archetypal dynamics and clustered by archetype (Figure 6C,D). In archetypes with differential dynamics across time, we find that transcripts that are prototypical of these archetypes are involved in the EGFR pathway (EGFR, RRAS), angiogenesis (RHOB, NOTCH1), endothelin signaling (END2, PIK3R1), inflammation (RRAS, SHC1), interleukin signaling (MKNK2, IRS1), and interferon *γ* signaling (ITGB1). We also observe differences between infected and bystander cells: EGFR, NOTCH1, RRAS, IRS1, PIK3R1, ITGB1, and SHC1 exhibit increased expression at later times in bystander cells while RHOB and NFKB2 exhibit increased expression at later times in infected cells. In both populations, MKNK2 and EDN2 expression decrease over time. In genes of particular interest to COVID-19, TMPRSS2, IL1RN, and IL1A have temporal dynamics most prototypical of archetype 1 (decreasing expression over time in both infected and bystander cells) while IFITM3 is most prototypical of archetype 3 (increasing expression in infected cells over time) (Figure 6D). Thus, CONDENSE used on gene dynamics across 3 days post-infection in HBECs highlights the divergence of infected and bystander genes in key signaling pathways related to immune response and regulation.

**Figure 6:**
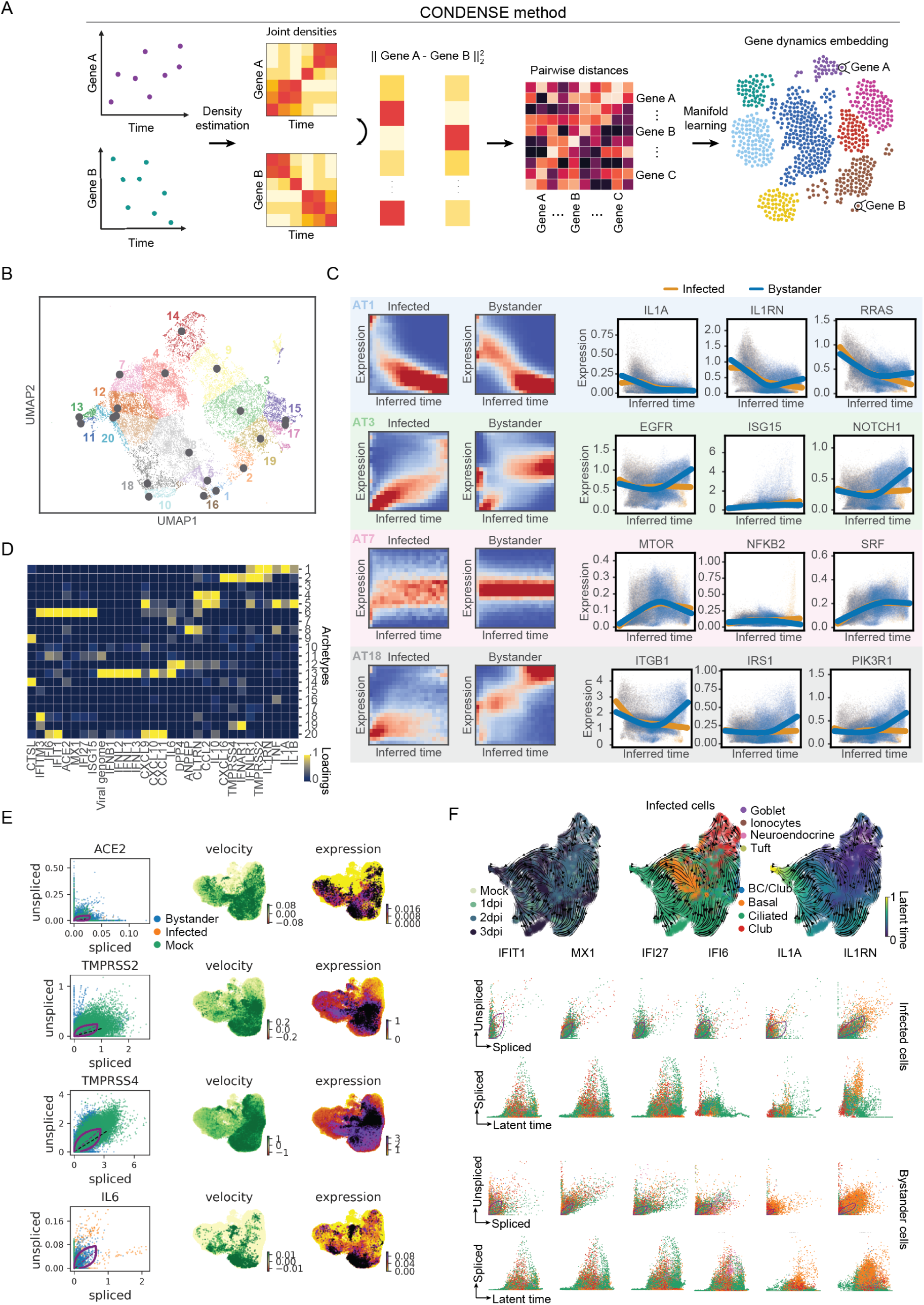
Transcriptomic dynamics across time in infected and bystander cells. (**A**) Overview of the CONditional DENSity Embedding (CONDENSE) method to characterize transcriptional dynamics over time (**B**) Clustering genes by their dynamical, joint density, stratified by infected and bystander cells, using archetypal analysis. (**C**) Images show conditional density estimates for archetypes identified in genes embedded into UMAP space. Scatter plots show LOESS fit for gene expression versus inferred times for various genes in archetypal clusters that exhibit differential dynamics across time. (**D**) Loading of viral and immune genes of interest onto the archetypes. (**E**) RNA velocity of genes of interest colored by infection status for individual cells. (**F**) RNA velocities fit separately with infected and bystander cells, highlighting genes of interest and latent time representations learned from RNA velocity.

### 2.7 Infection dynamics via RNA velocity

Next, to investigate transcriptional dynamics on the timescale of hours, we analyzed the RNA velocity of individual cells by fitting a dynamical model of transcriptional state based on spliced and unspliced transcript counts (Figure 6E) [27]. By embedding the RNA velocities in UMAP for all samples, we identified two distinct populations of infected cells - one in a high transition state, which corresponds to high velocity outflow streams, and another at a terminal state. Ciliated cells also exist in two RNA velocity states, one at an endpoint more transcriptionally different than infected cells, and another at a steady state more transcriptionally similar to infected cells. IL6 in infected cells is either in an induced state or repressed state, while few infected cells express IL6 at steady state (Figure 6F). ACE2 in bystander cells appear to be predominantly in an induced state while TMPRSS2 and TMPRSS4 have moderate RNA velocities, with the highest transcriptional changes occurring in basal or club cells. Using RNA velocity to compute a latent time validates that infected ciliated cells are driven to a terminal point, which also seems to mimic the transcriptional transition of infected basal cells (Figure 6F). Expression of IL1RN in both infected and bystander basal cells increases over time and exists predominantly in a steady state, while IFIT1, MX1, and IFI27 have high velocities in an induced transcriptional state in infected ciliated cells but a repressed state in bystander ciliated cells. In fitting separate models for infected and bystander cells, the most dynamical genes in infected cells are involved in inflammation (C5AR1, NFKBIA), interleukin signaling (IRS1), and EGFR signaling (EGFR, EREG). EGFR signaling and NFKBIA are induced early and are more highly expressed early in latent time while C5AR1 and IRS1 are more highly expressed at later time points in infected cells. In contrast, the most dynamica genes of bystander cells are involved in angiotensin II stimulated signaling (PRKCA), apoptosis (MAP3K5), EGFR signaling, and inflammation (ITGB1). In bystander cells, most of these genes are highly expressed at a later time. Collectively, RNA velocity validates that some of the most dynamic transcripts are involved in immune system regulation and highlights that genes identified as differentially expressed in the gene expression data are also associated with large RNA velocities in infected ciliated cells.

### 2.8 Cell tropism in clinical COVID-19 endotracheal aspirate

To compare our findings to an *in vivo* context, we performed scRNA-seq on endotracheal aspirates of a COVID-19 patient at the time of intubation (peak disease onset) and after extubation (during disease resolution), as well as on a control sample (see Methods). In these samples various immune cells, in addition to epithelial cells were identified (Figure 7A,B). Relative to the control patient, the COVID-19 patient had a higher abundance of neutrophils and monocytes (Figure S7A). Cells expressing at least one viral transcript from the endotracheal aspirate taken during intubation were identified but no infected cells were identified from the extubated sample, consistent with extubation following clinical recovery. In particular, ciliated cells were identified and found to express viral transcripts in the intubated endotracheal aspirate sample Figure 7C). However, alveolar type I cells were most highly infected by percentage of the identified cells with viral transcript (Figure 7C). Epithelial cells in the intubated sample show increased expression of immune system regulatory genes, in accordance with the *in vitro* HBEC data (Figure 7D,E,F,G). In addition, TMPRSS2, TMPRSS4, and ACE2 are expressed higher in the epithelial endotracheal aspirate cells of the intubated sample, relative to the extubated sample (Figure S7). However, IL1 and IL6 expression is higher in the extubated sample, relative to the intubated sample. Increased IL1 and IL6 expression in the extubated sample could be explained by the patient’s development of a secondary pneumonia. However, diagnosing secondary pneumonia in intubated and critically ill pediatric patients is imprecise. Nonetheless, the intubated patient was treated with antibiotics, potentially explaining the discrepancy between HBEC and clinical samples.

**Figure 7:**
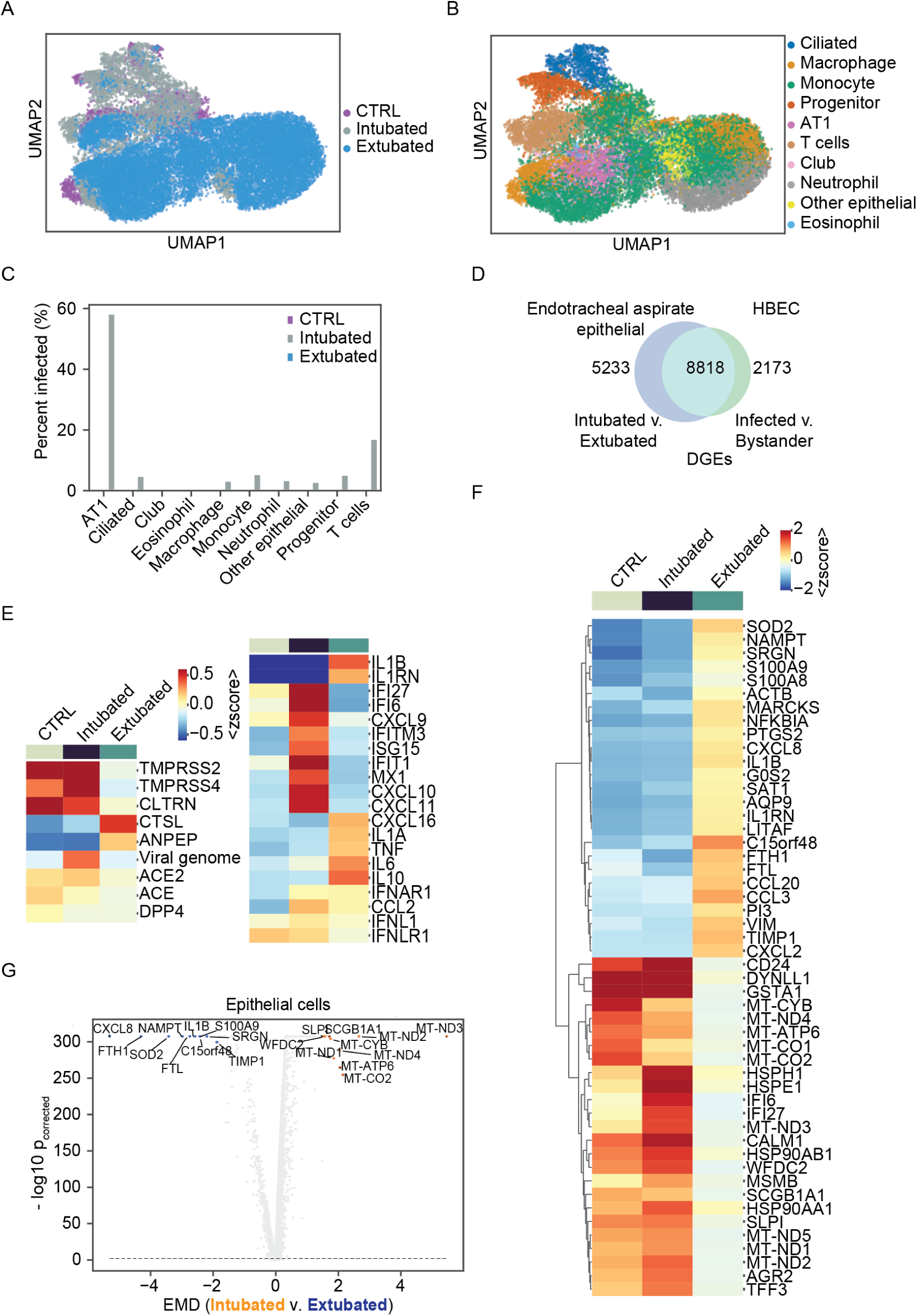
SARS-CoV-2 transcriptional dynamics in COVID-19 pediatric airway. (**A**) Single cells from endotracheal aspirate fluid from a healthy control and a pediatric patient with COVID-19 at the time of intubation and extubation. (**B**) Cell types identified in endotracheal samples. (**C**) Abundance of various cell types in each clinical sample. (**D**) Percentage of cells per cell type with at least one read aligned to the viral genome. (**E**) Expression of immune system genes of interest in epithelial cells from pediatric BALFs. (**F**) Expression of top 25 differentially expressed genes between intubated and extubated samples amongst epithelial cells from pediatric samples. (**G**) Differential gene expression for distributional distance versus corrected p-value for epithelial cells in pediatric endotracheal aspirate.

## 3 Discussion

To effectively treat COVID-19, we must first understand how SARS-CoV-2 causes disease and why the clinical presentation varies from asymptomatic infection to lethal disease. Here, we report the first longitudinal single-cell transcriptomic analysis of SARS-CoV-2 infected respiratory epithelium using an organoid model that reproduces the orientation key properties of the airway epithelium. The transcriptional data generated is of high quality, with an average of between 2,400 to 3,600 unique genes detected per condition. Our data reveals several novel viral transcripts and our methodology differentiates infected from bystander cells. Further, we demonstrate that ciliated cells are the major target cell of SARS-CoV-2 infection in the bronchial epithelium at the onset of infection and that cell tropism expands to basal, club, and BC/club cells over time. We also reveal that SARS-CoV-2 potently induces IFN in infected cells resulting in broad ISG expression in both infected and bystander cells. We also observe potent induction of the pro-inflammatory cytokine IL-6 and chemokines, which likely contribute to the inflammatory response *in vivo*. Furthermore, we find that differentially expressed genes between infected and bystander ciliated cells are enriched for age-associated genes, suggesting a potential mechanism for the increased severity of disease in elderly patients.

Single-cell transcriptomics enabled us to elucidate the SARS-CoV-2 transcriptome at single-cell resolution in multiple primary cell types over time. We developed a novel method to differentiate productively infected cell types by the distribution of the sub-genomic and genomic viral transcripts. We also identified polyadenylated viral transcripts remote from the 3’ end of the viral genome, which was unexpected given our sequencing method. Our RT-PCR validation experiments confirm the production of at least two unique, (Transcription regulatory sequence (TR)S-independent transcripts with poly-A tails that do not appear to result from non-specific oligo-d(T) priming. As the reported recombination rate for coronaviruses is high [28, 29] it is possible these short reads correspond to non-specific polymerase jumping. However, recent studies have identified TRS-independent chimeric RNAs produced during SARS-CoV-2 infection of Vero cells, a small portion (1.5%) of which are fused in frame [30]. Taken together with our results, this may suggest non-canonical sub-genomic RNAs with coding potential are produced during SARS-CoV-2 infection; however, this would require further validation. HCoV-299E non-structural protein 8 (nsp8) was recently shown to possess template-independent adenyltransferase activity [31]. Because poly-A tails play important roles in the stability and translation potential of canonical SARS-CoV-2 sgRNAs, it is interesting to speculate that coronaviruses might rely on the production of non-canonical, poly-adenylated sgRNAs to serve as decoys for cellular deadenylases. This would result in preservation of the poly-A tails of the genomic and subgenomic RNAs. Indeed, the production of sgRNAs during flaviviral infections is important for resistance to cellular exoribonucleases and innate immune evasion [32, 33].

Identification of the cell types infected by SARS-CoV-2 informs pathogenesis. We find that SARS-CoV-2 primarily infects ciliated cells, basal, club, and BC/club cells. This may result in aberrant function of these critical cell types. Ciliated cells, which are abundant in the respiratory epithelium, propel mucus and associated foreign particles and microbes away from the lower airway. Our finding that ciliated cells are the predominant target cell of SARS-CoV-2 infection in the bronchial epithelium has several important implications. First, dysfunction of ciliated cells by infection by SARS-CoV-2 may impair mucociliary clearance and increase the likelihood of secondary infection. Second, asthma, chronic obstructive pulmonary disease, and smoking are associated with both cilia dysfunction and increased severity of COVID-19. Whether these underlying conditions alter ciliated cells and thus increase their susceptibility to infection remains unclear, though SARS-CoV-2 has been identified in the ciliated cells of hospitalized patients, highlighting a potential connection between ciliated cells’ susceptibility to SARS-CoV-2 infection and COVID-19 disease progression [34]. Third, gene expression changes induced by SARS-CoV-2 infection in ciliated cells mirror the natural changes in expression that occur with aging, potentially providing insight into why elderly patients experience worse disease outcomes. The cell tropism of SARS-CoV-2 in the nasal epithelium and lower airway remain important areas of future investigation which will further enhance our understanding of COVID-19 pathogenesis.

Disease in COVID-19 patients is characterized by a lag following transmission with symptom onset at day seven and disease severity peaking 14 days post infection [35, 36]. This is in contrast to seasonal human coronaviruses and implicates an important role for the host immune response in COVID-19 progression. Several recent studies have revealed the induction of innate immunity during SARS-CoV-2 infection is dependent on viral growth kinetics and multiplicity of infection. [37, 38]. Here, we show that the innate response to SARS-CoV-2 is intact and rapid, as characterized by interferon, chemokine, and IL-6 induction. Interestingly, while we do not observe broad depletion of virus-susceptible cell populations, we detect increased expression of cell-death associated genes, which suggests the host anti-viral response is cytotoxic and may contribute to disease pathogenesis. Consistent with this, IL-6 is a potent pro-inflammatory cytokine and serum IL-6 levels predict respiratory failure [39]. Therapies targeting the IL-6 receptor are currently in clinical trials for the treatment of COVID-19. This work raises a number of important future directions including whether other airway and endothelial tissues similarly interact with SARS-CoV-2 and how these interactions vary *in vitro*.

## 4 Methods

### 4.1 Air-liquid interface culture of HBECs

HBECs, from Lonza, were cultured in suspension in PneumaCult-Ex Plus Medium according to manufacturer instructions (StemCell Technologies, Cambridge, MA, USA). To generate air-liquid interface cultures, HBECs were plated on collagen-coated transwell inserts with a 0.4-micron pore size (Costar, Corning, Tewksbury, MA, USA) at 5*x*10^4^ cells/*ml* per filter and inserted into 24 well culture plates. Cells were maintained for the first 3 days in PneumaCult-Ex Plus Medium, then changed to PneumaCult-ALI Medium (StemCell Technologies) containing the ROCK inhibitor Y-27632 for 4 days. Fresh medium, 100 *µl* in the apical chamber and 500 *µl* in the basal chamber, was provided every day. At day 7, medium at the apical chambers were removed, while basal chambers were maintained with 500 *µl* of PneumaCult-ALI Medium. HBECs were maintained at air-liquid interface for 28 days allowing them to differentiate. Medium in the basal chamber was changed every 2-3 days (500 *µl*).

### 4.2 Viral infection

SARS-CoV-2 isolate USA-WA1/2020 was obtained from BEI reagent repository. All infection experiments were performed in a Biosafety Level 3 facility, licensed by the State of Connecticut and Yale University. Immediately prior to infection, the apical side of the HBEC ALI culture was gently rinsed three times with 200 *µl* of phosphate buffered saline without divalent cations (PBS-/-). 10^4^ plaque forming units (PFU) of SARS-CoV-2 in 100 *µl* total volume of PBS was added to the apical compartment. Cells were incubated at 37°*C* and 5% *CO*_2_ for 1 hour. Unbound virus was removed and cells were cultured with an air-liquid interface for up to three days. Infections were staggered by one day and all four samples were processed simultaneously for single-cell RNA sequencing, as described below.

### 4.3 Sample preparation for single-cell RNA sequencing

Inoculated HBECs were washed with 1X PBS-/- and trypsinized with TrypLE Express Enzyme (ThermoFisher, Waltham, MA, USA) to generate single-cell suspensions. 100*µl* of TrypLE was added on the apical chamber, incubated for 10 min at 37°*C* in a *CO*_2_ incubator, and was gently pipetted up and down to dissociate cells. Harvested cells were transferred in a sterile 1.5 ml tube and neutralized with DMEM containing 10 percent FBS. An additional 100 *µl* of TrypLE was placed on the apical chamber repeating the same procedure as above for a total of 30 min to maximize collection of cells. Cells were centrifuged at 300 x g for 3 min and resuspended in 100 *µl* DMEM with 10 percent FBS. Cell count and viability was determined using trypan blue dye exclusion on a Countess II (ThermoFisher Scientific). The targeted cell input was 10,000 cells per condition. The Chromium Next GEM (Gel Bead-In Emulsion) Single Cell 3’ Gel beads v3.1 kit (10X Genomics, Pleasanton, CA, USA) was used to create GEMs following manufacturer’s instruction. All samples and reagents were prepared and loaded into the chip and ran in the Chromium Controller for GEM generation and barcoding. GEMs generated were used for cDNA synthesis and library preparation using the Chromium Single Cell 3’ Library Kit v3.1 (10X Genomics) following the manufacturer’s instruction. Generated libraries were sequenced on NovaSeq 6000 system using HiSeq 100 base pair reads and dual indexing.Cells were sequenced to an average depth of 31,383 reads per cell. The human genome, Ensembl GRCh38.98.gtf, and the SARS-CoV-2 genome, NCBI Genome database accession MT020880.1, were combined and used for alignment. We ran the standard 10X Genomics cellranger pipeline with a combined human and SARS-CoV-2 genome to obtain count matrices for each of the 4 growth conditions. Per condition, there were an average of between 10,000 to 15,000 counts per cell or an average of 2,400 to 3,600 unique genes detected per condition. Library preparations and sequencing were performed by the Yale Center for Genome Analysis.

### 4.4 Quantitative RT-PCR of SARS-CoV-2

Viral RNA from SARS-CoV-2 infected HBEC cell lysates was extracted using TRIzol (Life Technologies) and purified using Direct-zol RNA MiniPrep Plus according to manufacturer’s instructions (Zymo Research, Irvine, CA, USA). A two-step cDNA synthesis with 5 *µl* RNA, random hexamer, and ImProm-II Reverse Transcriptase (Promega, Madison, WI, USA) was performed. The qPCR analysis was performed in duplicate for each of the samples and standard curves generated using SARS-CoV-2 nucleocapsid (N1) specific oligonucleotides from Integrated DNA Technologies (Coralville, IA, USA): Probe: 5’ 6FAM ACCCCGCATTACGTTTGGTGGACC-BHQ1 3’; Forward primer: 5’ GACCCCAAAATCAGCGAAAT 3’; Reverse primer: 5’ TCTGGTTACTGCCAGTTGAATCTG 3’. The limit of detection was 10 SARS-CoV-2 genome copies/*µl*. The virus copy numbers were quantified using a control plasmid which contain the complete nucleocapsid gene from SARS-CoV-2.

### 4.5 Validation of polyadenylated SARS-CoV-2 transcripts

Huh7.5 cells grown in DMEM containing 10 percent FBS were infected with 10^4^ PFU of SARS-CoV-2 and cell lysates were harvested at 0, 1, 2, and 3 dpi. Using 0.3 *µg* total RNA extracted from mock or SARS-CoV-2-infected Huh7.5 cells at different time points, reverse transcription was performed with oligo-d(T)20 (ThermoFisher) and MarathonRT, a highly processive group II intron-encoded RT. MarathonRT purification and RT reactions were performed as previously described [40]. PCR (NEBNext Ultra II Q5 ® Master Mix, NEB, Ipswich, MA, USA) was performed with a gene-specific forward primer designed 700-nt upstream of the apparent boundary between the SARS-CoV-2 genome body and the putative poly-A tail. Oligo-d(T)20 was used as a reverse primer. Touchdown PCR cycling was used to enhance specificity of the PCR reaction. RT-PCR products were resolved on a 1.3% agarose gel with ladder (100 bp DNA Ladder, 1 kb Plus DNA Ladder, Invitrogen). Forward PCR oligonucleotides used in this experiment are below, which includes two positive controls.

Primer Name Position on Genome 5’-3’ Sequence:

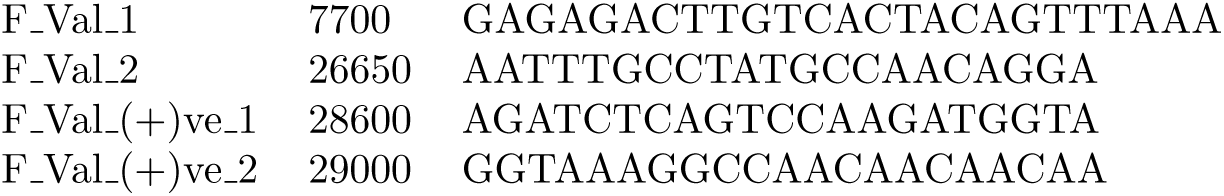

### 4.6 Electron microscopy

The cells were fixed using 2.5% glutaraldehyde in 0.1 M phosphate buffer, osmicated in 1% osmium tetroxide, and dehydrated in increasing ethanol concentrations. During dehydration, 1% uranyl acetate was added to the 70% ethanol to enhance ultrastructural membrane contrast. After dehydration the cells were embedded in Durcupan. 70 nm ultrathin sections were cut on a Leica ultramicrotome, collected on Formvar coated single-slot grids, and analyzed with a Tecnai 12 Biotwin electron microscope (FEI).

### 4.7 Cytokine measurement by multiplex immunoassay

Levels of IL6, IL1A, IL1B, IL1RN, and CXCL9 in the basolateral supernatants of mock and infected HBECs were all performed by EVE technologies (Calgary, AB, Canada) using the multiplex immunoassay analyzed with a BioPlex 200. All statistical analysis was performed using Prism GraphPad version 8. All were statistically analyzed using nonparametric Kruskal–Wallis test are indicated with a bar and the p-value is represented by a symbol. (* p-value ¡0.05, ** p-value ¡0.01, ***p¡0.001).

### 4.8 Clinical samples

#### Ethical statement

Human subjects research was performed after approval from the Yale University Institutional Review Board and conducted according to the principles expressed in the Declaration of Helsinki. Written informed consent was obtained from the legal guardian prior to sample collection.

#### Patient history and clinical sample collection

A 15 year old male with obesity, obstructive sleep apnea, and metabolic syndrome was presented with 6 days of fever (107°*F*), headache, cough, chest tightness/pain, fatigue, and anorexia. The patient then developed shortness of breath 2 days prior to admission and required escalating respiratory support and was intubated on day 2 of his hospital course. He was also diagnosed with hypoxic respiratory failure and acute respiratory distress syndrome (ARDS) secondary to COVID-19 infection. The patient remained intubated for 8 days. At the time of intubation, he had an oxygenation index of 28 and his chest radiograph demonstrated multi-focal bilateral airspace opacities. Laboratory chemistry were remarkable for lymphopenia with absolute lymphocyte count of 0.5 x 1000/*µL*, elevated ferritin (2386 ng/mL), elevated d-dimer (30.7 mg/L), elevated CRP (298 mg/L), transaminitis (ALT 245 U/L, AST 119 U/L), elevated BNP (835 pg/mL), elevated IL-6 (37 pg/mL), slightly elevated IL-10 (20 pg/mL), and slightly elevated IL-2R (1203 pg/mL). On the fifth day of being in mechanical ventilator, the patient developed secondary bacterial pneunomia and was treated with antibiotics. The control patient is a 15 year old female who was intubated for elective orthopedic surgery in early 2019. She arrived to the pediatric intensive care unit intubated for pain control and did not have any lung disease. She was extubated several hours after arrival once a pain control regimen was established. Endotracheal aspirates were collected at time of intubation and extubation. Approximately 5 ml of endotracheal aspirates were obtained and placed on ice. The endotracheal aspirates were passed through a 100 *µm* nylon strainer to remove debris and clumps, centrifuged and resuspended in cold DMEM with 10

### 4.9 scRNA-seq data analysis

#### 4.9.1 Cell type annotation

We used the standard scRNA-seq analysis pipeline for clustering [41]. Briefly, to account for transcript dropout inherent to scRNA-seq, we removed genes that were expressed in fewer than 3 cells and removed cells that expressed fewer than 200 genes. Next, we filter out cells with more than 10 percent of mitochondrial genes. We did not find a correlation between viral copy number and mitochondrial expression. The resulting raw unique molecular identifier (UMI) counts in each cell were normalized to their library size. Then, normalized counts were square-root transformed, which is similar to a log transform but does not require addition of a pseudo count. Data pre-processing was performed in Python (version 3.7.4) using Scanpy (version 1.4.6) [42].

We visually observed batch effects between conditions in 2-dimensional cellular embeddings. To remove these batch effects for clustering, cell-type annotation, and visualization, we used an approximate batch-balanced kNN graph for manifold learning (BB-kNN batch-effect correction) using Scanpy’s fast approximation implementation [43, 42]. BB-kNN assumes that at least some cell types are shared across batches and that differences between batches for a same cell type are lower than differences between cells of different types within a batch. For each cell, the 3-nearest neighboring cells in each condition were identified by Euclidean distance in 100-dimensional Principal Component Analysis (PCA) space. This kNN graph was used as the basis for downstream analysis.

To visualize the scRNA-seq data we implemented various non-linear dimension reduction methods and used the BB-kNN batch-corrected connectivity matrix as input for Uniform Manifold Approximation and Projection (UMAP) [44] and Potential of Heat-diffusion for Affinity-based Trajectory Embedding (PHATE) [45]. UMAP projections were generated using a minimum distance of 0.5. PHATE projections were generated with a gamma parameter of 1.

For cell clustering we used the Louvain community detection method [46] with the BB-kNN graph. We used high-resolution community detection and merged clusters based on expression of bronchial epithelium cell-type markers in order to isolate some rare cell types, e.g. Tuft cells [47, 48]. To annotate the different cell types present in HBECs we analyzed expression of a range of marker genes that were reported in a molecular cell atlas from Travaglini et al. [47]. We focused on 8 cell types: (i) Basal cells (KRT5, DAPL1, TP63), (ii) Ciliated cells (FOXJ1, CCDC153, CCDC113, MLF1, LZTFL1), (iii) Club cells (SCGB1A1, KRT15, CYP2F2, LYPD2, CBR2), (iv) BC/club (KRT4, KRT13), (v) Neuroendocrine cells (CHG1, ASCL1), (vi) Tuft cells (POU2F3, AVIL, GNAT3, TRPM5), (vi) Ionocytes (FOXI1, CFTR, ASCL3) and (viii) goblet cells (MUC5AC, MUC5B, GP2, SPDEF).

To analyze the ∼ hours-timescale splicing dynamics and transcriptomic regulation within infected and bystander cells undergoing changes across days of exposure to SARS-CoV-2, we aligned reads to segregate spliced and unspliced counts [49]. From these layered counts matrices, we compute RNA velocity for individual cells using scVelo, a generalization of the original RNA velocity method that allows for characterization of multiple transcriptional states [27, 49]. We calculate the first and second order moments with 50 PCs and assume 30 neighbors. Then, we fit three dynamical RNA velocity models in order to avoid the confounding bias of batch effects and to learn separate latent times from computed RNA velocities: one model including all HBECs, and one model each for infected and bystander cells. We visualize the learned dynamics and the relation of spliced and unspliced counts with scVelo and globally view the velocity in our original UMAP embedding.

#### 4.9.2 Infection threshold and infection score

Counting a viral transcript in a cell does not mean the cell is infected, as this count can come from virus attached to the surface of the cell, ambient virus in the suspension, or from read misalignment. Given the reported shared 3’ poly(A) tail in coronavirus transcripts [50], we were unsure whether we could correctly capture the different ORFs using the 10x Genomics 3’ gene expression library. Therefore, we aligned the viral reads to a genome-wide single “exon,” i.e., a count is given for a read mapped to SARS-CoV-2 ORFs and intergenic regions. These counts were used to infer individual cells’ infectious state. To filter out cells with viral genome transcript counts that may result from viral-cell surface attachment, ambient virus in the droplet suspension, or read misalignment, we considered infected cells to have ≥ 10 viral transcripts counts. This value was determined by a threshold of viral counts in the mock condition. While the mock condition is not expected to have viral counts, we did observe a small number that we attribute to misalignment. We observed only 5 mock cells with full SARS-CoV-2 viral genome transcript counts ≥ 10 transcripts. These criteria allowed us to find 144 infected cells at 1 dpi, 1428 cells at 2 dpi and 3173 cells at 3 dpi.

To quantify the extent to which an individual cell is transcriptionally similar to an infected cell, we used a previously developed graph signal processing approach called Manifold Enhancement of Latent Dimensions (MELD) [51]. We encoded a raw experimental score for each cell in the dataset such that -1 represents a bystander or uninfected cell and +1 represents an infected cell. Using the kernel from the BB-kNN graph (described above), these raw scores were smoothed in the graph domain, yielding an “infection score” per cell that represents the extent to which an individual cell is transcriptionally similar to infected cells. For summary statistics, this score was stratified by cell type and condition.

#### 4.9.3 Viral genome read coverage analysis

To visualize the viral read coverage along the viral genome we used the 10X Genomics cellranger barcoded binary alignment map (BAM) files for every sample. We filtered the BAM files to only retain reads mapping to the viral genome using the *bedtools intersect* tool [52]. We converted the BAM files into sequence alignment map (SAM) files in order to filter out cells that were removed in our single cell data preprocessing pipeline. The sequencing depth for each base position was calculated using *samtools count*. To characterize read distribution along the viral genome we counted transcripts of 10 different ORFs: ORF1ab, Surface glycoprotein (S), ORF3a, Envelope protein (E), Membrane glycoprotein (M), ORF6, ORF7a, ORF8, Nucleocapsid phosphoprotein (N) and ORF10.

#### 4.9.4 Differential Expression Analysis

To find differentially expressed genes across conditions, we used a combination of three metrics: the Wasserstein or Earth Mover’s distance, an adjusted p-value from a two-sided Mann-Whitney U test with continuity and Benjamini-Hochberg correction, and the binary logarithm of fold change between mean counts. Significance was set to *p*_*adjusted*_ ≤ 0.01. The Earth Mover’s distance, or 1-dimensional Wasserstein distance can be defined as the minimal cost to transform of distribution to another, and was previously used to assess gene expression that significantly differ between conditions [53, 54]. We performed several binary comparisons for each timepoint and for pooling 1, 2, and 3 dpi: infected vs. bystander, infected vs. mock cells, and bystander vs. mock cells. The 30 most differentially expressed genes (up- or downregulated, ranked by Wasserstein distance) in each condition, cell type, and analysis were represented in heatmaps. To identify putative cellular functions changed across conditions, we performed PANTHER-GO [55] statistical over-representation tests for up-regulated genes in each cell type and condition separately, using the default Human PANTHER-GO reference list as a background.

### 4.10 CONditional DENSity Embedding (CONDENSE) for unbiased characterization of gene dynamics

To characterize genes based on their temporal dynamics, we developed a new computational method that combines ideas from graph signal processing, density estimation, information theory, and manifold learning. Our method, called ***CON****ditional* ***DENS****ity* ***E****mbedding* (or CONDENSE) is designed to infer temporal dynamics, estimate joint and conditional densities, and then embed genes using these densities. Figure 6A shows an overview of the CONDENSE method. The goal of CONDENSE is to obtain an embedding of genes according to their transcriptional dynamics over time so that this embedding can be used for discovering new gene dynamics signatures.

#### 4.10.1 CONDENSE overview and notation

The CONDENSE algorithm for analyzing gene dynamics in single-cell data is as follows:

1. Obtain the condition of interest using pseudotime or other trajectory inference method
2. Obtain a joint density estimate over the cells for each gene as a function of time using kernel density estimation
3. Normalize the per gene density estimate by its temporal condition to obtain a conditional density embedding of genes
4. Use the gene dynamics embeddings for downstream analyses such as visualization, clustering, archetypal analysis, and gene-set enrichment

While the method was designed with single-cell expression data in mind, CONDENSE can be applied to any dataset of *m* examples 𝕏 = { **x**^(1)^, *…*, **x**^(*m*)^ } with a condition *c*. Here, we treat each gene as a sample, with expression of that gene across the scRNA-seq dataset represented by a vector **x**^(*i*)^ for a particular gene *i*. We infer a pseudotime to represent the condition *c*. Lastly, to compare the differential gene dynamics between strata of interest, *S*_*k*_ for *k* ∈ { 1, 2, *… l*}, we compute a conditional density embedding for each stratum separately and then concatenate the estimates to derive a joint distribution,

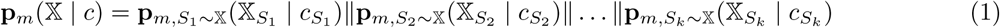

where **p**_*m*_(𝕏 | *c*) is the conditional density embedding, represented as a vector, for gene *m*. Across all genes, we get a conditional density embedding matrix, 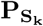 over *k* strata, which has a conditionally normalized joint probability density estimate for *m* genes. From this collection of conditionally normalized joint probability distributions, we perform manifold learning for an unsupervised view of gene dynamics. By contrasting gene dynamics for our two strata of interest, namely, infected v. bystander cells, we effectively compute a Wasserstein distance (or Earth Mover’s Distance, EMD) between conditionally normalized joint probability distributions, 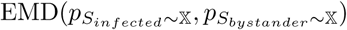 [56]. Next, we describe in detail each aspect of CONDENSE.

#### 4.10.2 Inferring a pseudotime condition

To infer a temporal condition *c* on an individual cell basis, constrained by a particular cell’s transcriptome, we used pseudotime or trajectory inference. Several pseudotime inference methods exist; however, it is not always clear what underlying dynamical process is captured by the latent or pseudotime representation [57]. To ensure that our condition corresponds to the dynamics over days post-infection rather than across other forms of cellular heterogeneity, we used a supervised approach to infer a pseudotime by smoothing a cells’ known timepoint label onto a continuum according to the transciptomic similarity between cells using Manifold Enhancement of Latent Dimensions (MELD) [51]. In our implementation, we used an *α*-decaying kernel with distances and connectivities derived from an approximate BB-kNN graph, allowing us to define a similarity matrix,

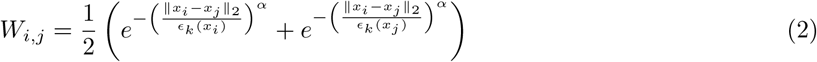

where *ϵ*_*k*_(*x*_*i*_) and *ϵ*_*k*_(*x*_*j*_) are the Euclidean distances in PCA space between cells *x*_*i*_ and *x*_*j*_ for the *k* neighbors of *i* with non-zero connectivities. We can obtain the total variation of a given signal **f** ∈ ℝ^*n*^ for our similarity graph **W** over *n* vertices via the Laplacian quadratic form,

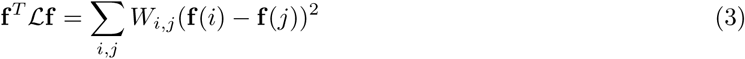

where ℒ is the graph Laplacian and **f** (*i*) is the timepoint label for a given cell pair *i*. Here, we assign **f** for a given cell *i* as **f**_*i*_ = *t*_*i*_ for *t* ∈ {0, 1, 2, 3}, indicating whether the cell was derived from the Mock, 1, 2, or 3 dpi sample. Thus, the graph signal we filter corresponds to the timepoint information drawn from the experimental design. Eigendecomposition of the graph Laplcian, ℒ = ΨΛΨ^−1^ yields eigenvectors 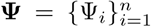 with eigenvalues that are local minima of **f** ^*T*^ ℒ**f**, enabling definition of the graph Fourier transform of the label per cell, 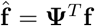 [51]. We use signal processing in the spectral domain with Laplacian regularization to obtain a filter matix,

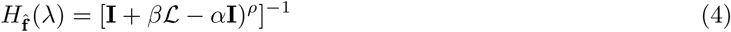

where *λ* are the eigenvalues of **Ψ**, *α* is a modulation parameter (set to 40), *β* is a reconstruction penalty (set to 1), and *ρ* specifies the order of the filter (set to 2). To obtain pseudotime, we follow the implementation of [51], which computes the frequency response or smoothed signal by using Chebyshev polynomial approximation to obtain *H* [58].

#### 4.10.3 Conditional density estimate

We next compute a conditional density estimate for each strata and each gene’s expression as a function of pseudotime. First, we compute a normalized density of each *m*-th gene’s expression across cells in a stratum of interest *S*_*k*_ with a kernel density estimator,

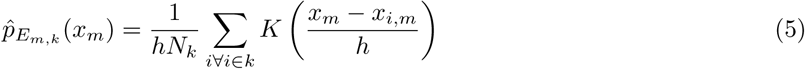

where *h* is the bandwidth of kernel *K*, 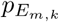 is the probability density for the *k*-th stratum of cells across the 95 percentile range of expression in that stratum, *N*_*k*_ is the number of cells in that stratum, and *x*_*i,m*_ is the normalized counts for gene *m* in cell *i*. Here, we use a uniform kernel function with a bandwidth *h* = 1. Similarly, we can estimate the probability density of inferred pseudotime across a stratum of cells, using the same kernel, *K*, to approximate 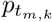. Now, to estimate the conditional density, we take the histogram, i.e., an un-normalized kernel density estimate, using a modified kernel *K*′ = *N*_*k*_ · *K*, and normalize a gene’s expression histogram for each stratum of interest by its pseudotime, yielding a conditional density estimate for each gene *m* in a stratum *k* of interest,

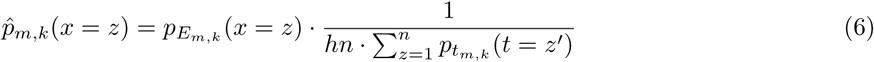

where *x* = *z* specifies the density estimate for cells at a binned expression interval *z*, and *t* = *z*′ specifies the cells at a binned condition or pseudotime interval *z*′. Here, we assume that *z* and *z*′ are intervals of fixed width such that 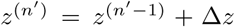 where 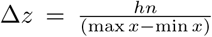 and 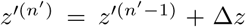 where 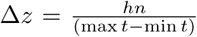 where *n*′ is an indexing parameter ∈ 1, 2, *…l* up to *n* and, in practice, we constrain the range of *x* and *t* to the 95-th percentile range. Thus, we obtain a conditional desnsity estimate for each gene and strata after iterating through each binned expression and conditional interval.

#### 4.10.4 Conditional Density Embedding

In practice, Equation 6 can be computed quickly using a two-dimensional histogram of gene expression versus pseudotime and normalizing each bin by the sum over the gene expression distribution per binned pseudotime interval. Here, we use 20 evenly spaced bins over the 95 − *th* percentile range of gene expression and pseudotime. We iterate this process for each gene and flatten the conditionally normalized, two-dimensional histogram (conditional density estimate) **P**_**m**,**k**_ into a vector **p**_**m**,**k**_. Then, we concatenate the vectors for each stratum according to Equation 1 to get a matrix of conditional density estimates across *m* genes, 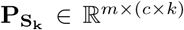, where the *m*-th row and *c*-th column species the conditional density estimate for gene *m* at pseudotime *c* in a particular stratum of interest, repeated *k* times. Thus, we obtain our conditional density embedding across *k* strata of interest, 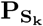.

#### 4.10.5 Archetypal analysis of CONDENSE matrix

To find extremal dynamics on an individual gene basis between strata, we performed archetypal analysis on the conditional density estimate matrix, 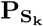. Archetypal analysis estimates the principal convex hull of a *d*-simplex dataset [59]. The convex hull of *d* dimensions can be thought of as an envelope around the data. We implement archetypal analysis by defining the principal convex hull as the (*d* − 1)-dimensional convex hull that best accounts for the data according to,

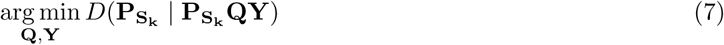

where *D* is a measure of distortion and **Q** and **Y** are the optimal matrices that minimize the distortion *D* subject to the constraints, |**q**_*d*_ |_1_= 1 and **Q** ≥ 0, and |**y**_*m*_ |_1_= 1 and **Y** ≥ 0, respectively. These constraints yield archetypes,

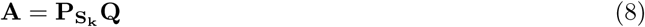

which are convex combinations that can be approximated by a weighted average of the feature vectors 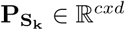 [60]. In this implementation of principal convex hull analysis (PCHA), we identify archetypes **A** that favor features representing “corners” of the data, which allows one to identify *archetypes* or distinct and extremal aspects of the data [60]. To estimate the *d*-dimensionality of our conditional density estimate data, we used the “elbow method” of cumulative variance explained by principal components, after computing 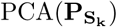 to obtain (*d* − 1) archetypes. Archetypal analysis is one of many low-rank matrix approximation methods, one of which is the more broadly used principal component analysis.

#### 4.10.6 Biological insights from CONDENSE

After identifying distinct gene dynamics by archetypal analysis, we further characterize the set of genes expressed in our dataset by visualizing the multidimensional, conditional density estimates in a single dimensionally-reduced embedding. Any manifold learning or dimesionality-reduction method is possible but we use the popular UMAP algorithm to learn a mapping *u* : ℝ^*c×k*^ ↦ ℝ^2^ [44]. We fit this function on the conditional density embedding, 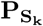 and transform 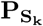 to get a conditional density embedding, 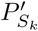 in UMAP space. In order to characterize the dynamics according to archetypal or extremal dynamics within the HBEC dataset, we assign each gene to an archetype by minimizing the Euclidean distance of each gene’s conditional density embedded point to the nearest archetypal point in UMAP space,

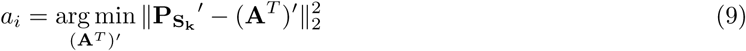

where *a*_*i*_ represents the archetype that satisfies the minimization problem, (**A**^*T*^) represents the archetypes in UMAP coordinates and 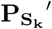 denotes the conditional density embedding of genes per strata *k* in UMAP coordinates. This allows us to identify genes with distinct and differential dynamics in infected versus bystander cells over the course of infection and globally visualize the variation in gene dynamics at single-gene resolution.

We internally validate CONDENSE by locally estimated scatterplot smoothing (LOESS) [61]. Briefly, we visualize gene expression for interesting archetypes against the cells’ inferred pseudotime to confirm that the conditional density estimate matches the dynamics across pseudotime for each strata. This also provides a complementary way to assess the differential dynamics between strata, albeit assuming that changes in gene expression are locally linear across pseudotime. After obtaining CONDENSE, there are many downstream analyses that could be done, including using associating the archetypes and genes with other cell or gene metadata via supervised learning. However, to provide additional insight into the genes exhibiting distinct and differential dynamics between infected and bystander cells, we performed PANTHER-GO analysis on genes in various archetypal clusters and identified over-represented pathways in archetypes showing interesting dynamics.

## 5 Code and Data Availability

All differential gene expression analyses and their associated metrics are publicly available at the Van Dijk Lab GitHub: https://github.com/vandijklab. Software for the CONDENSE method and scripts used to analyze the scRNA-seq data are also available at the Van Dijk Lab GitHub: https://github.com/vandijklab. The annotated scRNA-seq data can be browsed with an interactive web-tool, courtesy of the Chan-Zuckerberg Initiative. The raw scRNA-seq data will also be made publicly available and deposited in NCBI’s GEO database.

## 6 Acknowledgments

We thank the Yale Center for Genome Analysis for help with sequencing and alignment. This work was supported by NIH grants K08 AI128043 (CBW), R21AI133440 (AW), R01AI141609 (AW), and K08AI119139 (EEF). CBW was also supported by a Burroughs Wellcome Fund Career Award for Medical Scientists and Robert Leet and Clara Guthrie Patterson Trust Award. CBW and DvD were supported by a Fast Grant (Emergent Ventures). AI and AMP are Investigators of the Howard Hughes Medical Institute.

## Author contributions

conceptualization, MMA, JW, NGR, VG, EFF, DvD, CBW; methodology, MMA, NGR, EFF, DvD, CBW; computational analysis NGR, VG, DvD; validation, MMA, CBW; formal analysis NGR, VG, DvD; investigation, MMA, RBF, JW, CBW; resources, BW, EFF; writing—original Draft, MMA, NGR, VG, DvD, CW; writing—review & editing, all authors; visualization, NGR, VG, DvD; supervision, DvD, CBW; funding acquisition, DvD, CBW.

---

**Figure S1:**
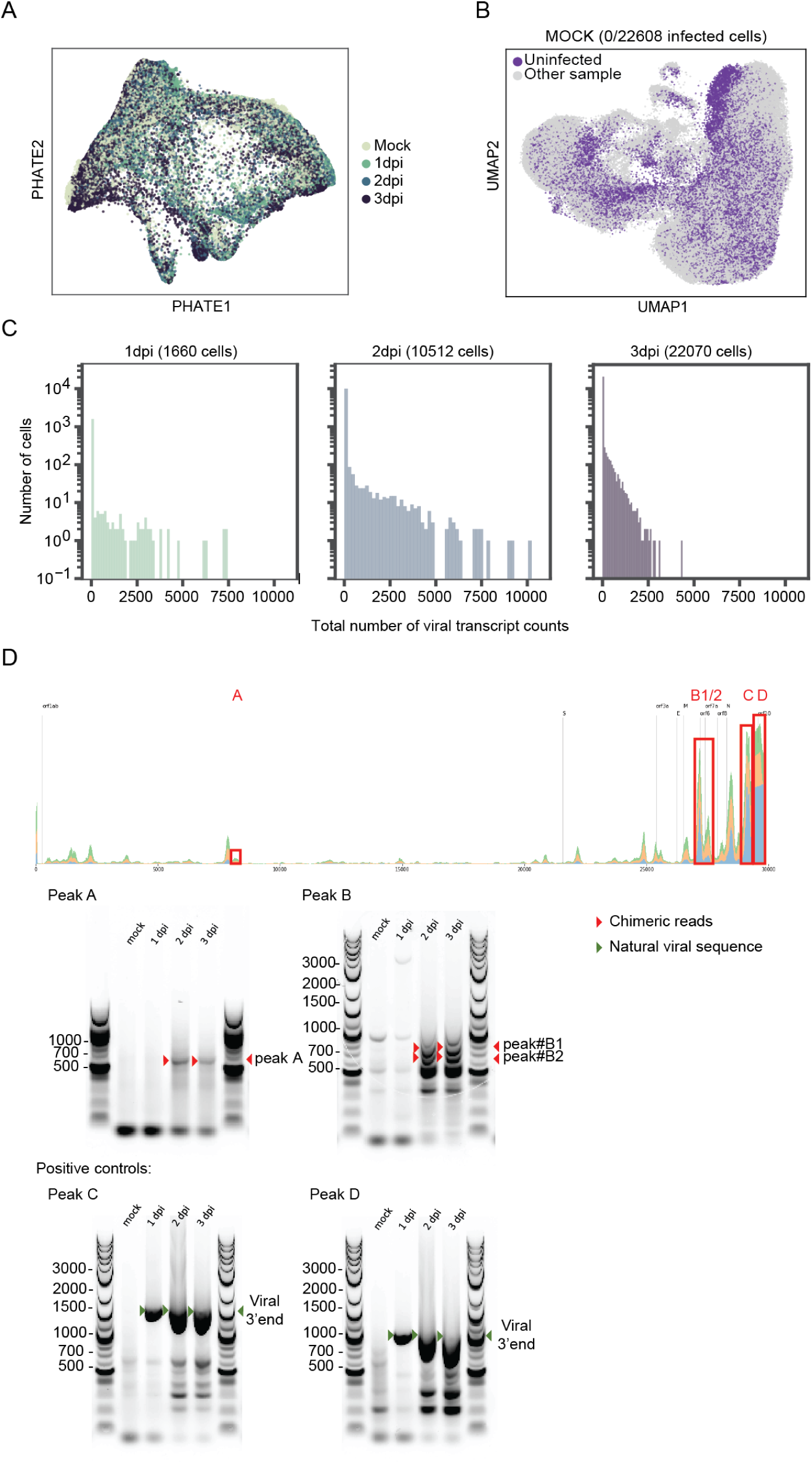
SARS-CoV-2 viral genome transcript counting to determine infections state. **A.** PHATE visualization of the scRNA-seq gene counts after batch correction. Each point represents a cell. Cells were colored according to their samples. **B.** UMAP visualization of the mock sample, 1 dpi, 2 dpi and 3 dpi are represented in Figure 1. **C.** Histograms of viral transcript counts per cell on a logarithmic scale for each condition. **D. Top panel**: Schematic of reads aligning across the SARS-CoV-2 genome. Red boxes indicate mapped reads validated with RT-PCR, including two previously unreported poly-adenylated transcripts (A, B). **Bottom panel**: RT-PCR spanning the junctions between poly-A tails and SARS-CoV2 genome body for non-canonical transcripts (Peaks A, B) and two positive controls (Peaks C, D). The products were run on agarose gels. Red arrowheads denote the expected amplicons for novel transcripts, while green arrowheads denote amplicons for the natural viral 3’end.

**Figure S2:**
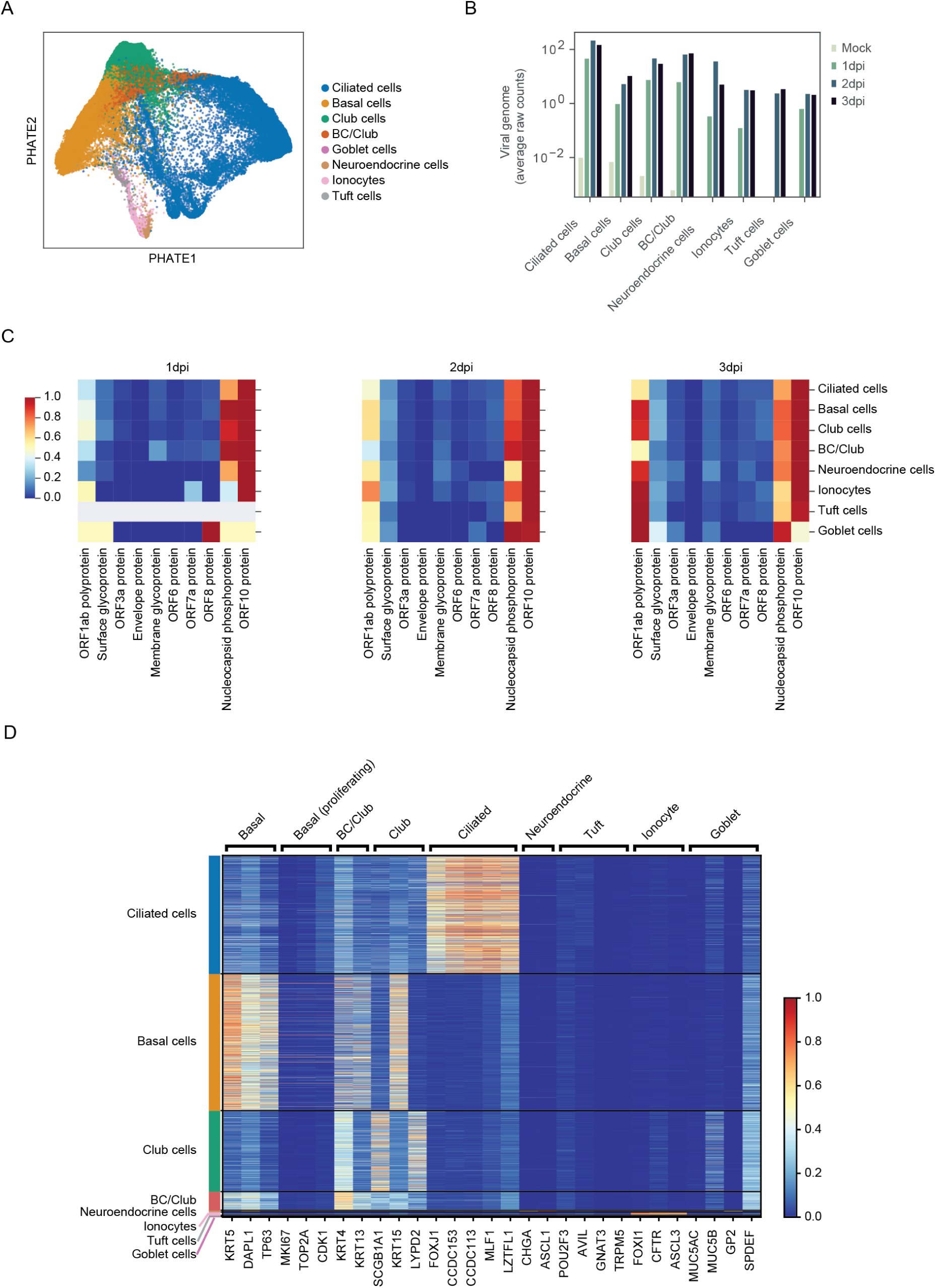
SARS-CoV-2 cell tropism and SARS-CoV2 sub-genomic expression across bronchial epithelial cell types. **A.** PHATE visualization of the cell types. **B.** Histogram of the average raw counts of viral transcripts per cell type across conditions. **C.** Normalized heatmap of the viral Open Reading Frame (ORF) counts in each cell type across three conditions: 1 dpi, 2 dpi and 3 dpi. **D.** Heatmap displaying expression of marker genes for each cell types and SARS-CoV-2 putative relevant genes.

**Figure S3:**
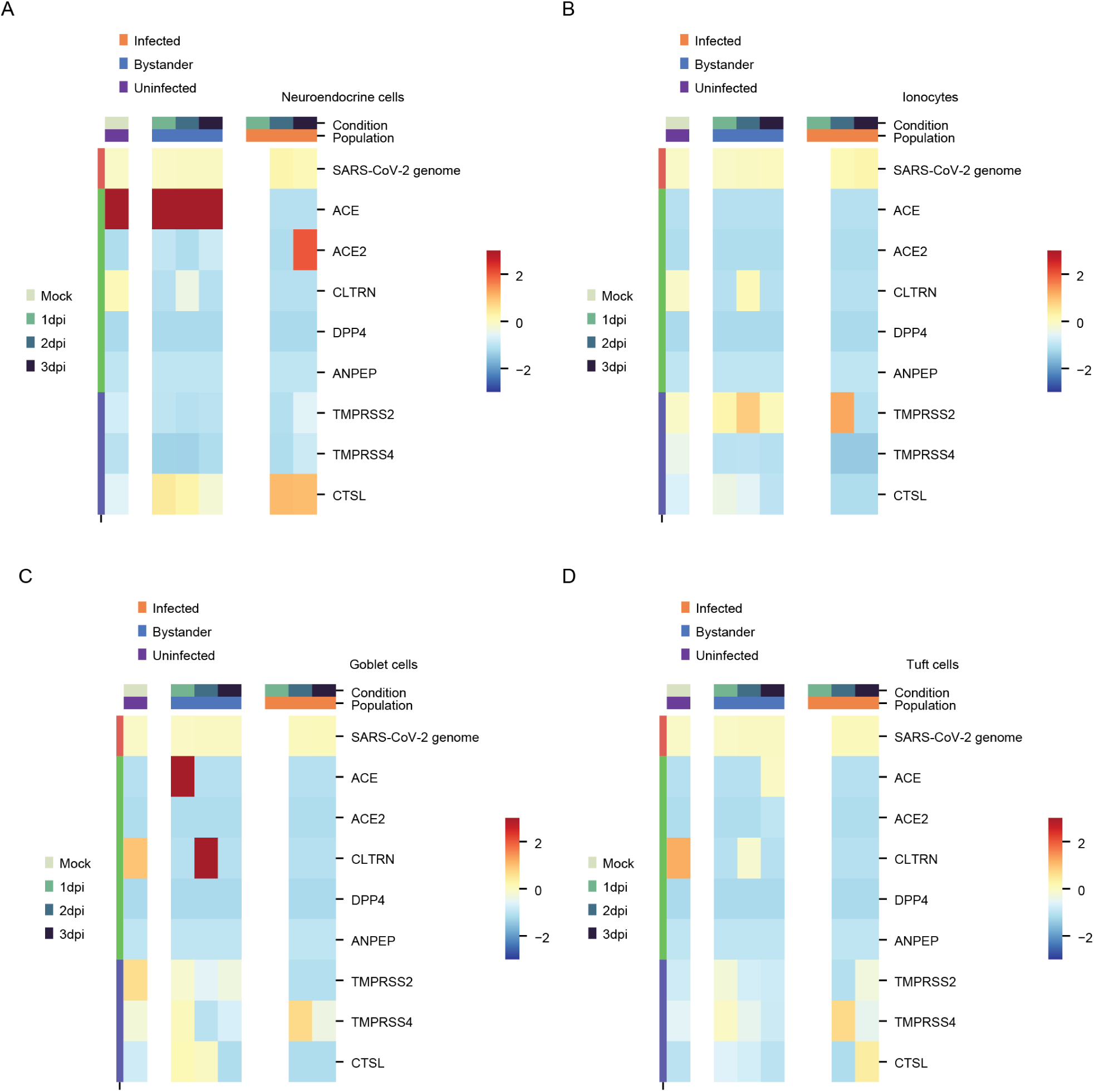
Expression of known and potential viral entry determinants across bronchial epithelial cell types. **A-D.** Heatmaps of receptors and proteases in neuroendocrine cells cells (**A.**), ionocytes (**B.**), goblet cells (**C.**) and tuft cells (**D.**).

**Figure S4:**
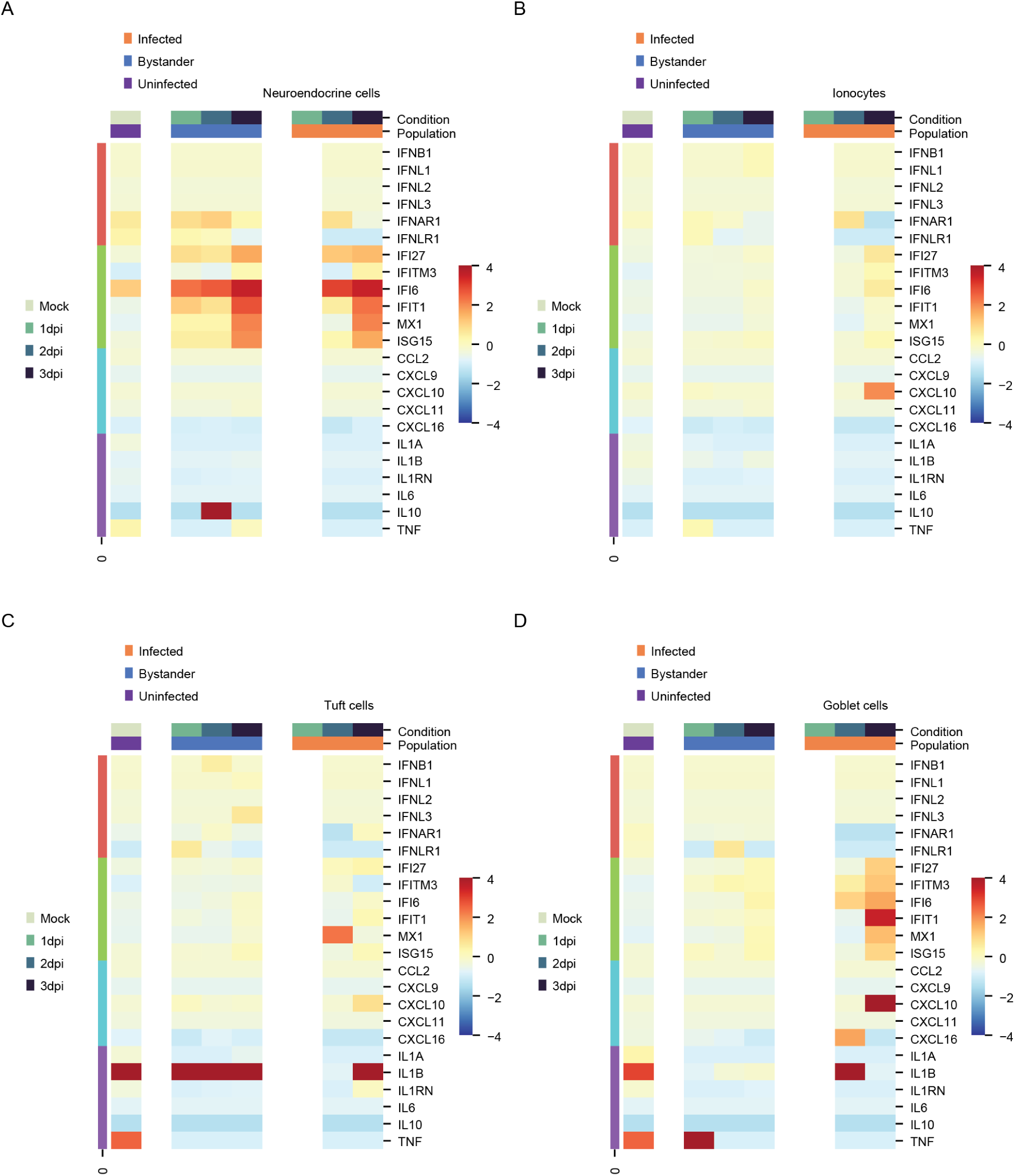
Innate immunity markers in SARS-CoV-2 infection. **A-D.** Heatmaps of cytokines, chemokines, interferons and interferon-stimulated genes in neuroendocrine (**A**), ionocytes (**B**), goblet (**C**) and tuft cells (**D**).

**Figure S5:**
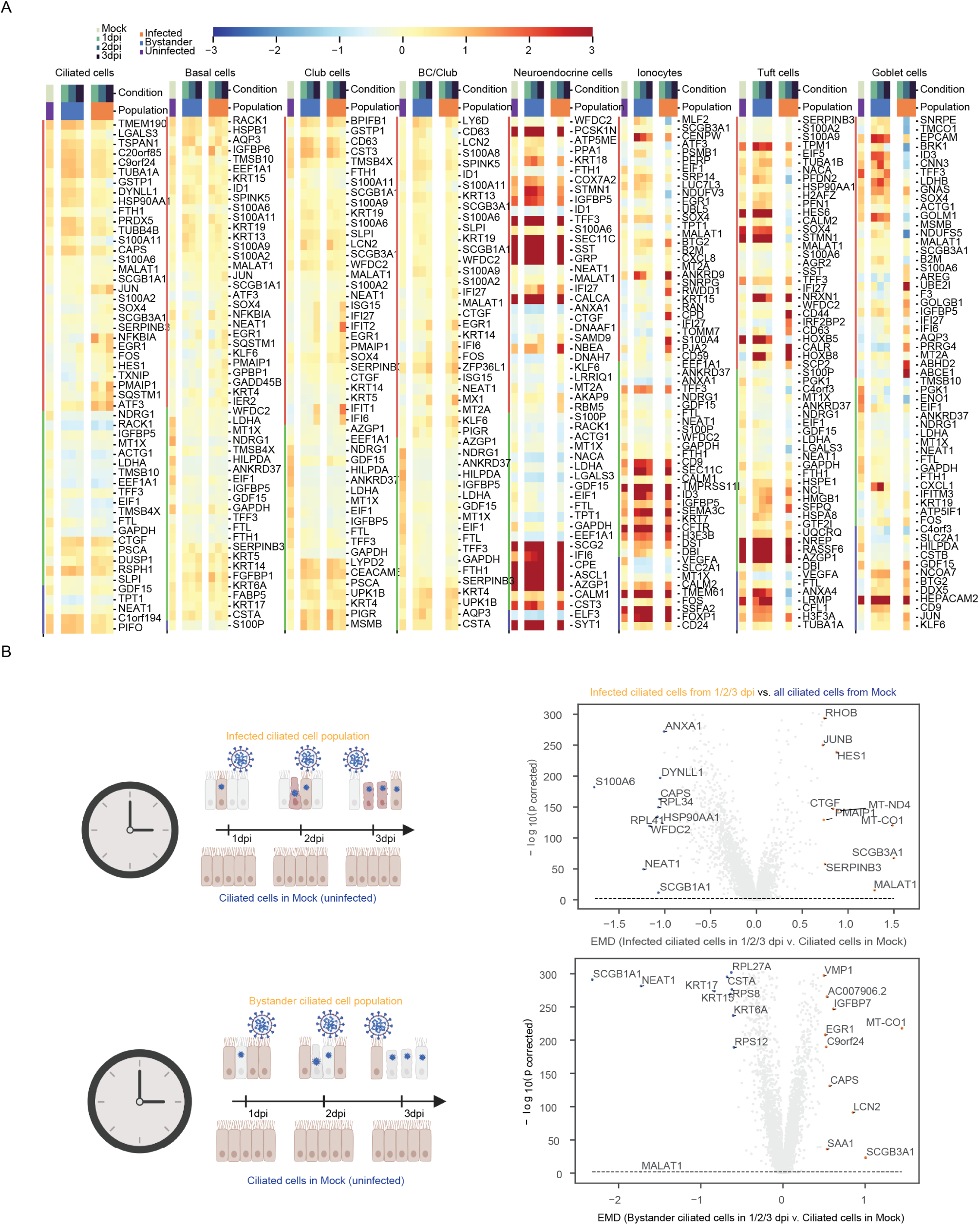
Expression of differentially expressed genes. **A.** Normalized heatmaps of the 30 most differentially expressed genes between uninfected, bystander and uninfected cells across cell types and conditions. Left to right : basal cells, BC/Club cells, club cells, neuroendocrine cells, ionocytes, goblet cells and tuft cells. **B.** Volcano plots highlighting the most differentially expressed genes between bystander and uninfected cells in ciliated cells.

**Figure S6:**
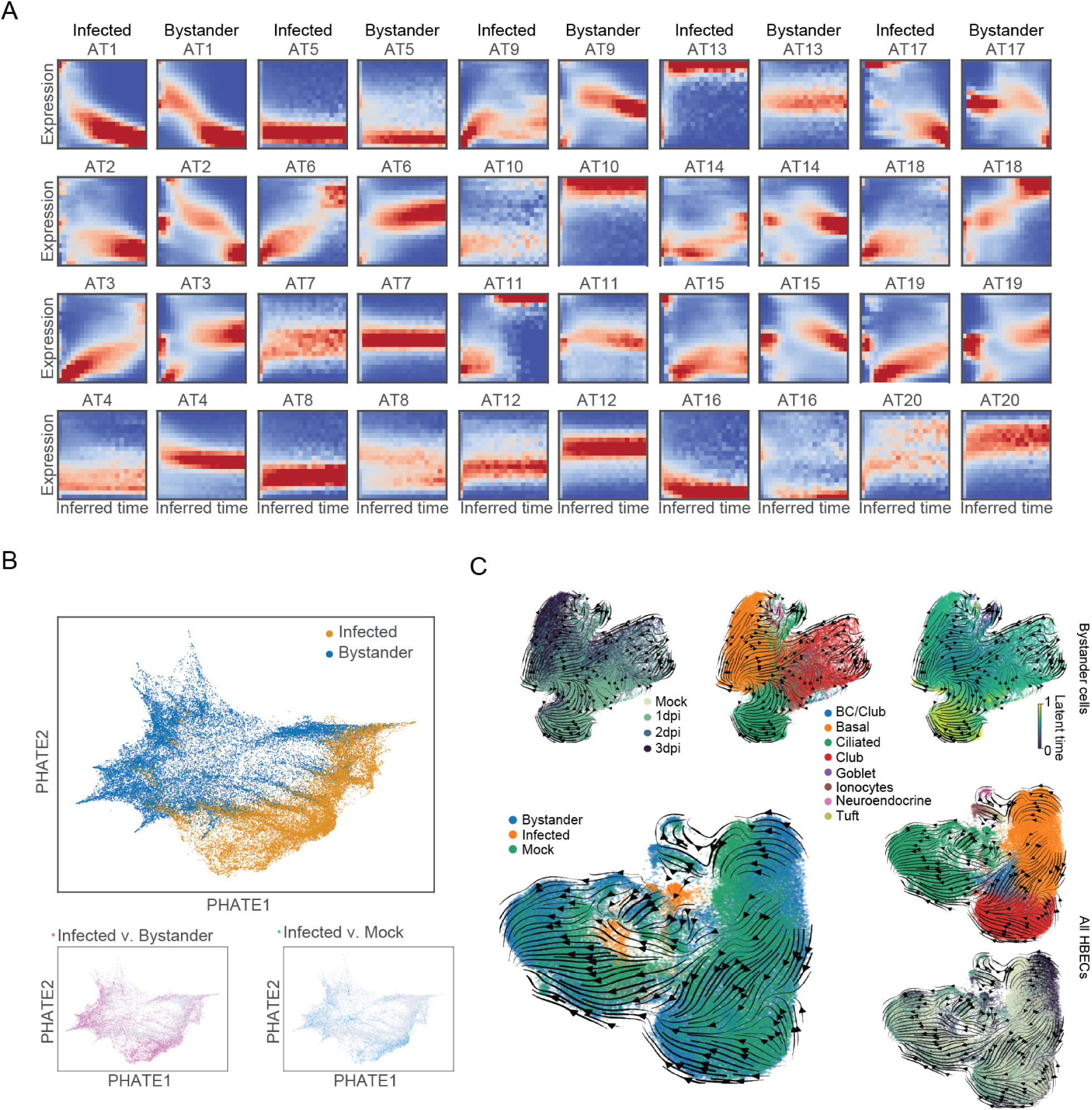
Archetypal gene dynamics and RNA velocity in bystander cells. (**A**) Archetypes derived from conditional density of gene expression versus time, stratified by infected and bystander cells. (**B**) Unsupervised manifold learning of conditional density images generated by separating infected and bystander per gene, highlighting the differential gene dynamics in infected and bystander cells for different differentially expressed genes (smaller scatter plots). (**C**) RNA velocity streams for bystander cells, stratified by timepoint, cell type, and latent time learned from RNA velocity (top). Velocity of genes of interest from dynamical RNA velocity model fit on all HBECs.

**Figure S7:**
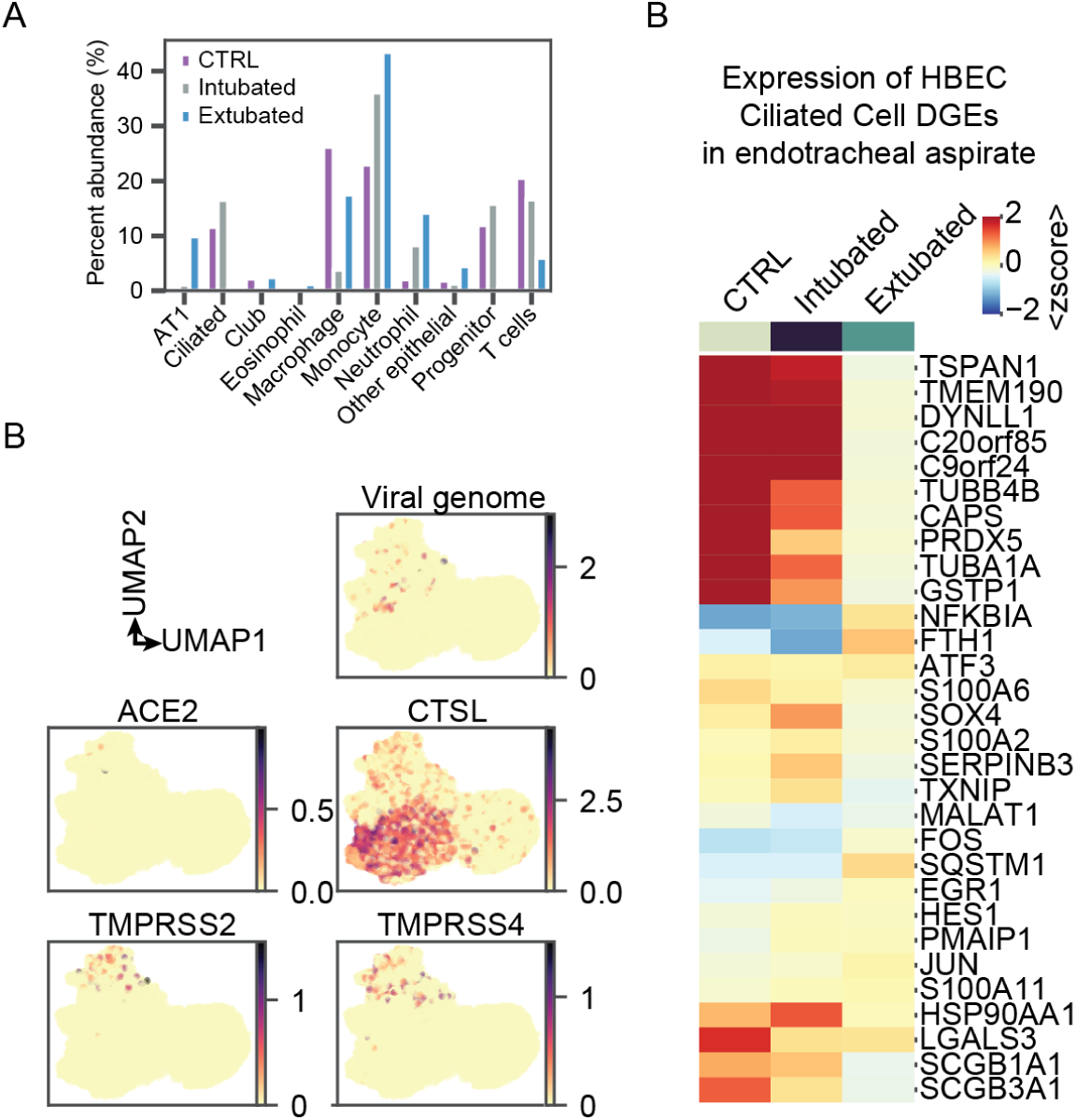
Expression of genes of interest in COVID-19 pediatric endotracheal aspirates. (**A**) Expression of SARS-CoV-2 genes of interest in epithelial cells (heatmap) and across all cells in the endotracheal aspirates (scatter plots). (**B**) Expression in epithelial cells from endotracheal aspirates of differentially expressed genes (infected v. bystander) in ciliated cells from the HBEC samples.

